# Structural analysis of HERC2/UBE3A and HERC2/DOCK10 complexes provides new insights into the molecular basis of Angelman, Angelman-like and Dup15q Syndromes

**DOI:** 10.1101/2025.09.16.670041

**Authors:** Auguste Demenge, JiaWen Lim, Arpoudamarie Roc, Boglarka Zambo, Gergo Gogl, Alexandra Cousido-Siah, Alastair Mc Ewen, Anna Bonhoure, Camille Kostmann, Christine Fagotto-Kaufmann, Pascal Eberling, Judit Osz-Papai, Alberto Podjarny, Eduardo Howard, Christelle Golzio, Anne Debant, Susanne Schmidt, Gilles Travé

**Affiliations:** Equipe Labellisée Ligue 2015, Département de Biologie Structurale Intégrative, Institut de Génétique et de Biologie Moléculaire et Cellulaire (IGBMC), INSERM U1258/CNRS UMR 7104/Université de Strasbourg, France; Institut de Génétique et de Biologie Moléculaire et Cellulaire (IGBMC), INSERM U1258/CNRS UMR 7104/Université de Strasbourg, France; Centre de Recherche en Biologie Cellulaire de Montpellier (CRBM), Université de Montpellier, CNRS, Montpellier, France; Facultad Regional Tierra del Fuego, Universidad Tecnologica Nacional, Ushuaia, Tierra del Fuego, Argentina

**Author notes:** equal contributors (Auguste Demenge, JiaWen Lim).

## Abstract

UBE3A and HERC2 are two mutually interacting HECT E3 Ubiquitin ligases whose genes are altered in 15q11.2-13.1 Duplication (Dup15q) Syndrome, Angelman Syndrome (AS) and other phenotypically-related mental retardation syndromes. Using quantitative binding assays, X-ray crystallography and sequence conservation analysis, we show that the HERC2/UBE3A complex occurs in probably most animals with a central nervous system, *via* a conserved interface involving the RLD2 domain of HERC2 and a “DxDKDxD” motif of UBE3A. We found that HERC2 also recognizes and binds to similar DxDKDxD motifs within a handful of other proteins relevant to brain development (DOCK10, PCM1, USP35, BAZ2B, ARID4A, ARIP4, RERE and MYT1). We further investigated the interaction of HERC2 with DOCK10, a RAC1- and CDC42-GEF protein that regulates dendritic spine morphogenesis in hippocampal neurons. Both disruption of the HERC2-binding motif in DOCK10 and knockdown of HERC2 affected the GEF activity of DOCK10. We also show that the DOCK10-induced dendritic spine formation is dependent on its ability to bind HERC2. Structural modeling of full-length DOCK10, free or bound to either RLD2, CDC42 or RAC1 indicates that the GEF activation of full-length DOCK10 requires a conformational change that is stimulated by binding to HERC2. Based on our data, we propose that under pathological conditions, in developing brains with an abnormal dosage of either HERC2 or its dominant partner UBE3A, increased or decreased amounts of HERC2/DOCK10 complexes could lead to altered GTPase activation. This in turn could affect dendritic spine formation and neurodevelopment.

## Introduction

Ubiquitin-protein ligase E3A (UBE3A, also called E6AP) is an E3 ubiquitin (Ub) ligase whose maternal allele is deleted, mutated or inactivated by *de novo* genetic alterations in the severe neurodevelopmental disease Angelman syndrome (AS) [1]. The maternal *UBE3A* allele is also duplicated or triplicated in Dup15q syndrome, a genetic condition that accounts for approximately 1-3% of ASD cases [2]. The specific involvement of the maternal *UBE3A* allele in neurodevelopmental syndromes is explained by the epigenetic silencing of the paternal allele of the *UBE3A* gene in most brain regions. However, both alleles are expressed elsewhere, allowing the paternal allele to compensate for the defective maternal allele in the rest of the body. This epigenetic mechanism is directly linked to the presence of the “Angelman Syndrome – Prader Willi Syndrome imprinting center” (AS-PWS-IC), proximal to the *UBE3A*-encoding gene [3, 4].

HERC2 is another E3 Ub ligase whose gene is also altered in rare neurodevelopmental syndromes, presenting several symptoms reminiscent of AS [5–8]. Remarkably, *UBE3A* and *HERC2* are both located in the 15q11q13 chromosomal region. Some of the systematic *UBE3A* maternal allele deletions observed in AS patients also involve the deletion of the neighboring *HERC2*. Furthermore, on the protein level, UBE3A and HERC2 belong to the same family of E3 Ub ligases. Both UBE3A and HERC2 contain a C-terminal HECT (Homologous to E6AP C-terminus) domain responsible for the ligase activity. The high sequence similarity of both domains suggests that the two genes originated from a duplication event during evolution, explaining their proximal chromosomal location. Moreover, UBE3A and HERC2 proteins have been shown to interact with each other [9–12]. A recent transcriptional study suggested that UBE3A and HERC2 proteins may cooperate in Dup15q syndrome [13]. The UBE3A-HERC2 complex thus emerges as a critical actor in brain development, whose perturbation may lead to neurodevelopmental disorders.

Here, we defined and solved, by X-ray crystallography, the structure of the interface of the HERC2-UBE3A complex, involving the RCC1-like Domain 2 (RLD2) of HERC2 and a “DxDKDxD” sequence motif of UBE3A. The HERC2-UBE3A interface is conserved in most animals with a centralized nervous system (CNS), pointing to the putative importance of this complex for brain development. We further identified and validated by X-ray crystallography and affinity measurements an additional set of HERC2-binding proteins containing the DxDKDxD motif (DOCK10, PCM1, USP35, BAZ2B, ARID4A, ARIP4, RERE and MYT1), all of which are also relevant to brain development. This suggests the possibility that the RLD2 domain of HERC2 establishes a tightly coordinated network of competing interactions, the imbalance of which may lead to neurodevelopmental defects. Using biochemical and cellular assays, we then focused on the interaction of HERC2 with DOCK10, a RhoGEF protein involved in dendritic spine morphogenesis [14], and showed that HERC2 expression and binding to DOCK10 affects DOCK10 activities. Finally, we present a compelling structural model of the activation of DOCK10 GEF activity upon binding to HERC2, that is fully consistent with our experimental data.

## Results

### The interface of the UBE3A-HERC2 interaction boils down to a complex between the HERC2 RLD2 domain and a short-disordered peptide of UBE3A

The UBE3A protein (852, 873 or 875 residues depending on the splicing isoform) is characterized by an N-terminal AZUL zinc-binding domain and a C-terminal E3 Ubiquitin-ligase HECT domain, while HERC2 (4834 residues, one of the 50 largest human proteins) harbors three RLD domains and a C-terminal HECT domain (Figure 1A). The HERC2-UBE3A interaction was previously found to involve the RLD2 domain of HERC2 and residues 150-200 of UBE3A (numbering of the 852-residues UBE3A human isoform I) [10, 11, 15]. The large size of HERC2 makes it challenging to isolate for *in vitro* quantitative binding assays. To overcome this problem, we used our recently developed native holdup assay coupled to Western blotting [16]. This assay allowed us to measure the affinity dissociation constant (K_d_) of purified full-length UBE3A immobilized on resin against non-purified, endogenous HERC2 from lysates of human neuroblastoma SH-SY5Y cells (Figure 1B). Depletion of HERC2 from the lysate with increasing amounts of recombinant full-length UBE3A revealed a K_d_ of 0.56 ± 0.15 μM (Figure 2A and Supplementary Figure 1). Conversely, we measured by native holdup assay the binding affinity of resin-bound, purified RLD2 domain of HERC2 (residues 2941-3342) to full-length endogenous UBE3A (Figure 1C) from the same cell lysates, revealing a K_d_ < 0.1 μM (Figure 2B, supplementary Figure 7). To further define the minimal HERC2-binding region of UBE3A, we synthesized a series of peptides spanning UBE3A residues 150-200 (Listed in supplementary Table 1). Preliminary experiments pointed to UBE3A residues 150-174 (HTKEELKSLQAKDEDKDEDEKEKAA) as being the strongest RLD2-binding peptide. Native holdup assay (Figure 1D) revealed a K_d_ = 0.34 ± 0.09 μM for the binding of this UBE3A peptide to full-length endogenous HERC2 (Figure 2C, Supplementary Figure 2). We next used fluorescence polarization assay (Figure 1E) to examine the binding of the peptides to purified HERC2 RLD2 domain (Figure 2D-E, Supplementary Figure 3 and Supplementary Table 1). The competitive fluorescence polarization (cFP) confirmed UBE3A residues 150-174 as the strongest RLD-binding peptide (Figure 2D, K_d_ < 0.1 μM). A shorter peptide encompassing UBE3A residues 160-174 also displayed specific binding to RLD2, albeit with weaker affinity (Figure 2E, K_d_ = 6.85 ± 0.29 µM). Altogether, these quantitative affinity data demonstrate that the HERC2-UBE3A interaction can be pinpointed to a minimal interface complex between the HERC2 RLD2 domain and UBE3A residues 160-174.

**Figure 1.**
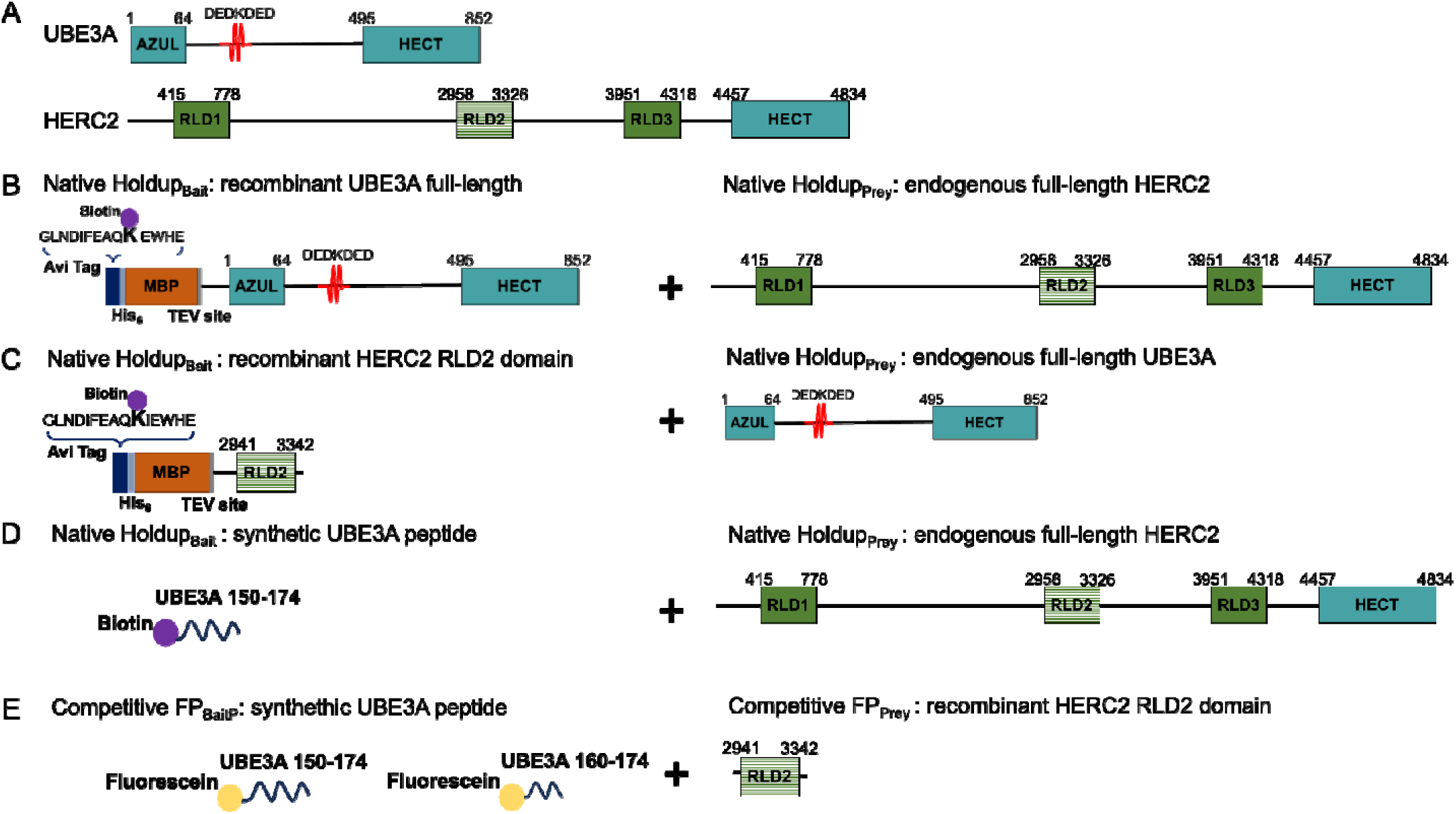
Schematic representation of the UBE3A and HERC2 E3 Ubiquitin-ligases and of derived constructs and peptides used in characterizing their interaction. A. Schematic diagram of full-length UBE3A isoform I (852 residues) and HERC2 (4834 residues). B-E. Schematic diagram showing the constructs of different biotinylated/fluorescein labeled-peptides or proteins with their binding partners used in native holdup and cFP assays.

**Figure 2.**
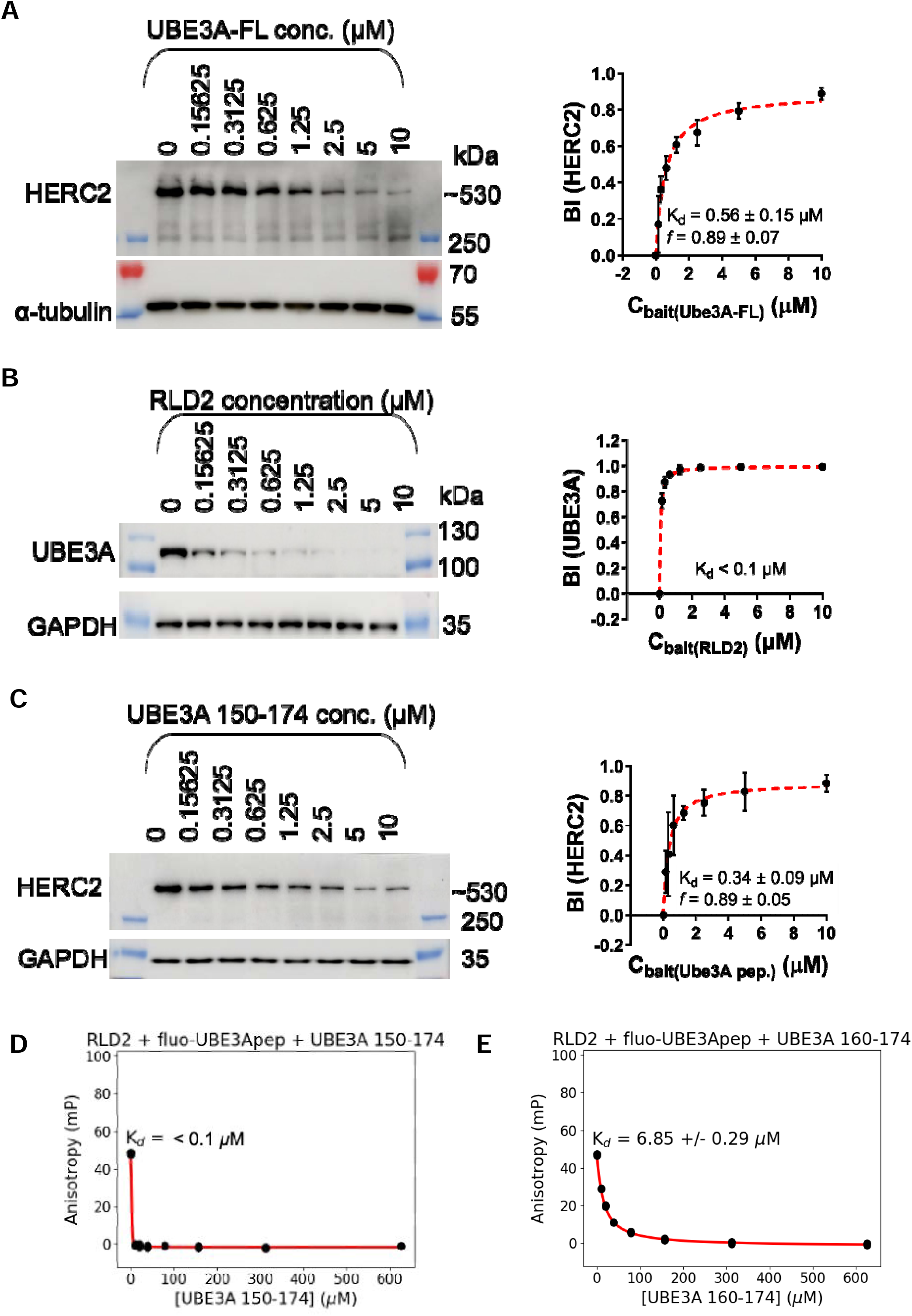
Determination of the binding affinity between full-length UBE3A or UBE3A 25-mer peptide with full-length HERC2 using native holdup titration. **A-C.** In the native holdup experiments (n=3), the streptavidin resins were saturated with biotinylated full-length MBP-fused UBE3A protein (A), MBP-fused HERC2 RLD2 domain (residues 2941-3342)(B) or UBE3A peptide containing amino acids 150-174 (HTKEELKSLQAKDEDKDEDEKEKAA)(C) and titrated from 10 μM to 0 μM to capture endogenous HERC2 or UBE3A protein from SH-SY5Y lysates. For holdup assays using MBP-fused proteins, streptavidin resin saturated with biotinylated MBP was used as a bait control, provided in increasing complementary amounts to the decreasing amounts of MBP-fusions. For holdup assays using biotinylated peptides, streptavidin resin saturated with biotin was used as the complementary bait control. The depletion of endogenous HERC2 from the liquid phase was followed by Western blot analysis. GAPDH or α-tubulin were used as a non-depleted control protein. Densitometry quantification of the protein bands was conducted with Fiji and a curve was fitted based on hyperbolic binding equation (red dashed line) to determine the binding affinity constant. **A.** Native holdup between full-length UBE3A and HERC2 from SH-SY5Y lysates. **B.** Native holdup between RLD2 domain and UBE3A from SH-SY5Y lysates. **C.** Native holdup between UBE3A 150-174 peptide and HERC2 from SH-SY5Y lysates. **D-E.** Binding affinities of the RLD2 to the different UBE3A peptides obtained by fluorescence polarization (FP). The reversibility of the complex formation was monitored with a competitive measurement by titrating the RLD2/fluo-UBE3Apep complex with an increasing amount of the different non-labeled UBE3A peptides as indicated. A decrease in the FP signal indicates the reversible complex formation. **D.** Competitive FP by UBE3A 25-mer peptide. **E.** Competitive FP by UBE3A 15-mer peptide. K_d_: equilibrium binding dissociation constant. Please see Supplementary Figure 1 and Supplementary Figure 2 for full blots of native holdup titrations; Supplementary Figure 3 and Supplementary Table 1 for the competitive FP by different UBE3A peptides.

### Crystal structure of RLD2 domain bound to UBE3A residues 160-174

Subsequently, we explored the crystallization of the RLD2 domain alone or in the presence of various peptides encompassing the UBE3A region 150-200. Crystals were readily obtained for isolated RLD2 as well as for RLD2 in the presence of the specific, yet weakly binding peptide comprising UBE3A residues 160-174. In contrast, despite repeated efforts, no crystals could be obtained for RLD2 in the presence of the longer and stronger-binding peptides comprising UBE3A residues 150-174 and 150-200. We hypothesized, that these longer peptides may contain two or more alternative RLD2-binding sites that could enhance RLD2-binding affinity by avidity effects, but they may also adopt alternative RLD2-binding poses, generating a structural heterogeneity that prevents crystallization. The crystallized complex consisting of HERC2 RLD2 (residues 2941-3342) bound to UBE3A residues 160-174 was solved at 3 Å resolution (Figure 3A). RLD2 displays a donut-shaped, seven-bladed β-propeller fold typical of the RCC1 family, with a central funnel connecting the “bottom” and “top” sides of the donut [17]. The UBE3A peptide lies on a highly conserved groove located on the top face of the RLD2 domain (Figure 3B). The seven HERC2-binding residues of UBE3A visible in the crystal are essentially charged, with a central lysine (Lys, K) flanked by aspartic (Asp, D) and glutamic acids (Glu, E) on both sides (sequence: DEDKDED).

**Figure 3.**
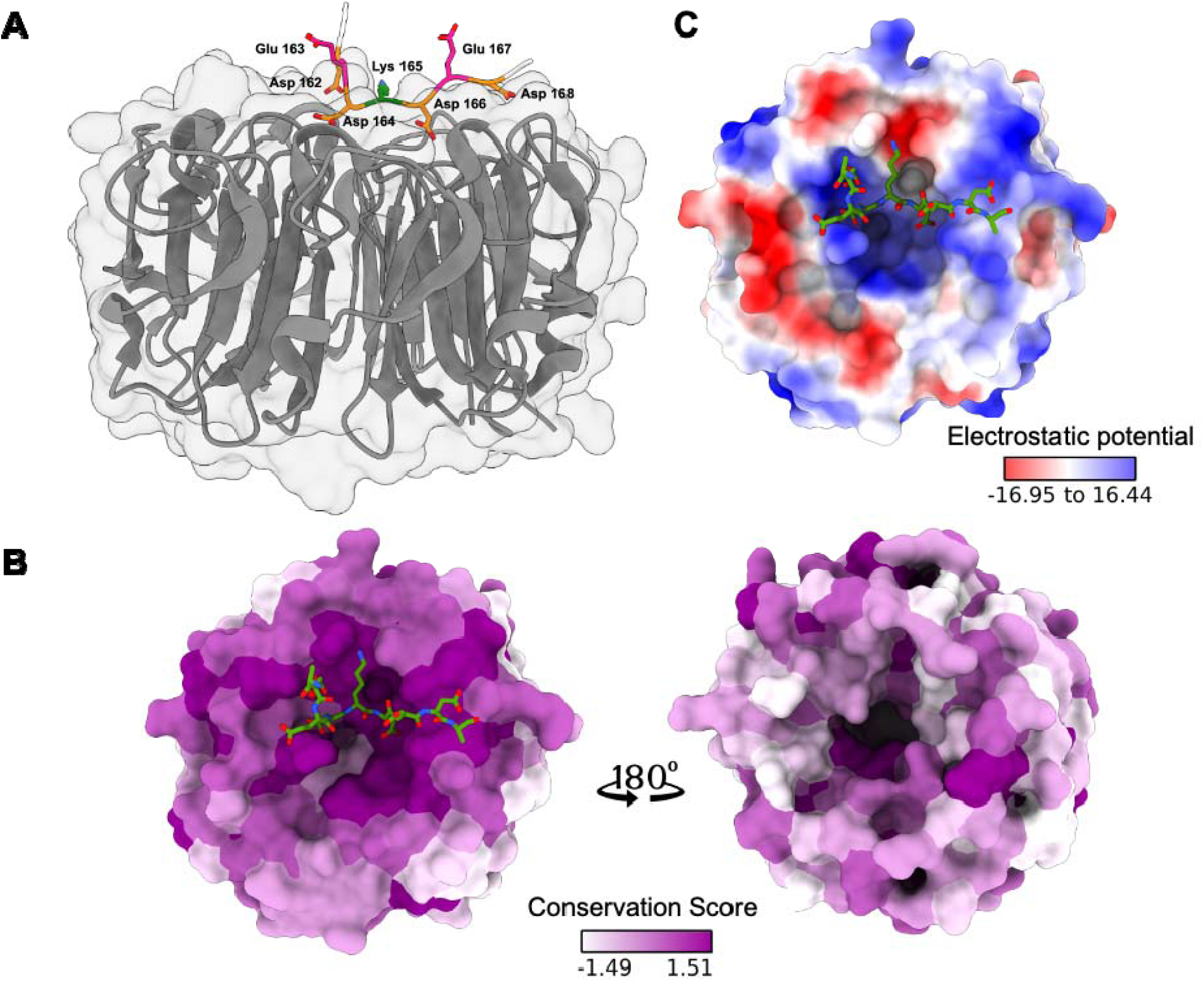
Crystal structure of HERC2 RLD2 domain bound to UBE3A 183-197 peptide. (PDB: 7Q41). **A.** RLD2 adopts the canonical donut-shaped, 7-bladed beta-propeller fold of RCC1-like domains. According to its conventional representation [17], the peptide interacts with the “top” side of the RCC1-like fold. The peptide is oriented from the N-terminus (left) to the C-terminus (right) in this representation and the following ones. **B.** AL2CO conservation score [65], calculated using the Chimera program from the HERC2 sequences of 9 representative animal organisms (see sequences in Figure 5A) and mapped on the surface of the RLD2 domain. Note the particularly high conservation (“hot” purple color) of the motif-binding groove. **C.** Surface electrostatic charges of the surface of the RLD2 domain, calculated using the Chimera program. A positively charged (blue) groove hosts the four aspartic acid residues (D) of the DEDKDED motif. The central lysine residue (K) is embedded in a negatively charged (red) pocket. Further details of the complex are shown in Figure 4. Please refer to Supplementary Table 2 for the data collection and refinement parameters.

Further visual inspection of the complex showed that the four Asp of the DEDKDED sequence are accommodated within a positively charged (blue) groove on the RLD2 surface (Figure 3C), establishing multiple hydrogen bonds (Figure 4A). Specifically, the first two Asp form hydrogen bonds with Ser 3160, Arg 3161, Gln 3215, Lys 3231 and Arg 3236 of RLD2. Notably, two water molecules are positioned at the interface and mediate some of these hydrogen-bonding interactions (Figures 4A and B). The central Lys of the DEDKDED sequence forms hydrogen bonds with Asp 3283, and Asp 3285 (Figure 4A and C), further stabilizing the interaction within a negatively charged (red) pocket on RLD2 (Figure 3C). Furthermore, the extended side chain of the central Lys also established hydrophobic interactions with Leu 3267 and Tyr 3234 (Figure 4A). Finally, the two C-terminal Asp of the DEDKDED sequence form hydrogen bonds with Ser 2978, Lys 2979, Ser 3001, Ser 3318, Ser 3319 and His 3286, thereby contributing further to the extensive electrostatic network at the interface (Figures 4A and D). In short, the central Lys and the four surrounding Asp of the DEDKDED sequence establish critical charged and hydrophobic interactions with the RLD2 domain. By contrast, the two Glu residues of that sequence point towards the solvent (Figure 3A) and are, therefore, not involved in the interface. The key HERC2-binding residues of UBE3A therefore boil down to the following sequence motif: DxDKDxD, where x stands for non-determining residues.

**Figure 4.**
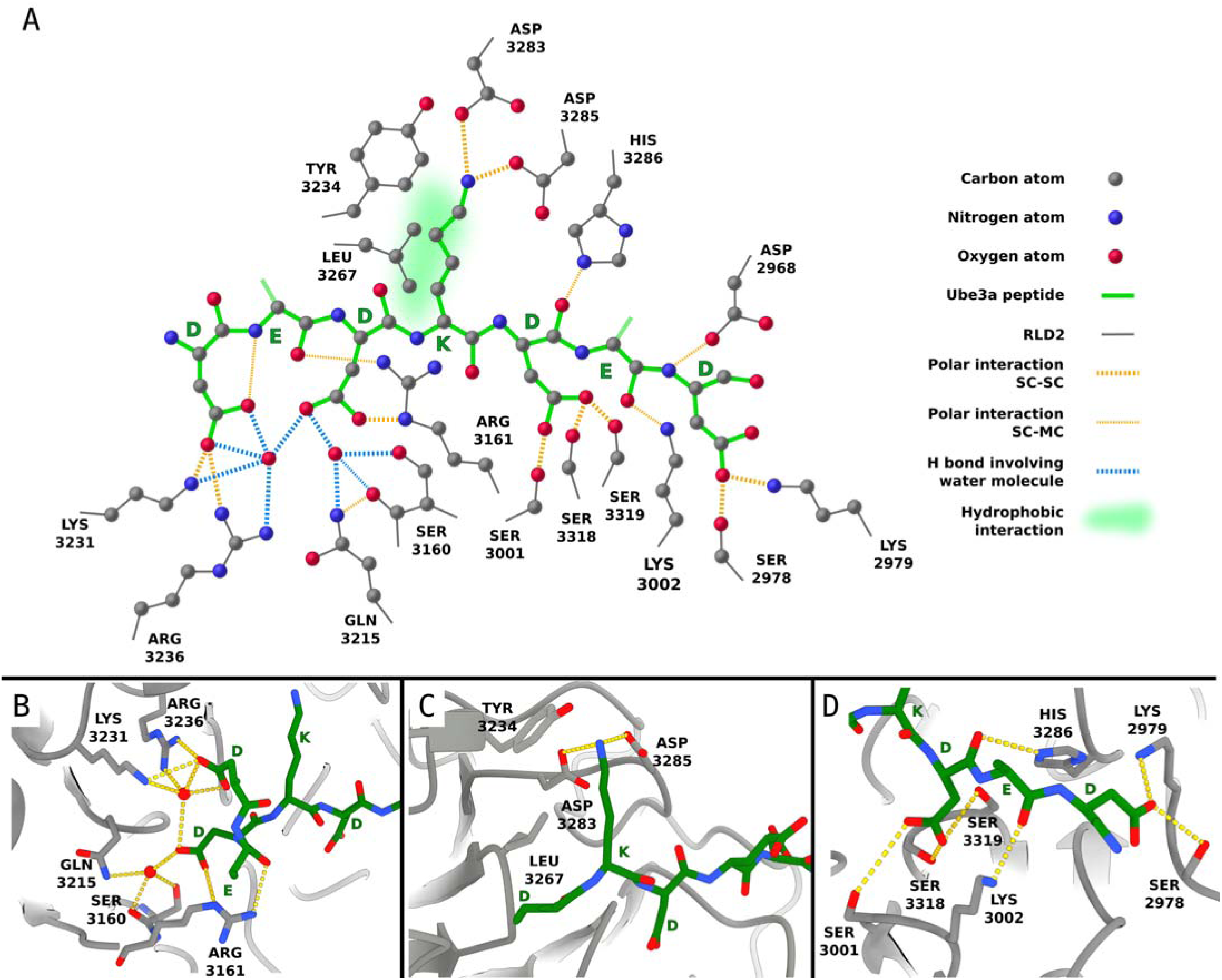
Interaction network between the DxDKDxD motif and the RLD2 domain. **A.** Schematized network of contacts between RLD2 domain and DEDKDED motif of the UBE3A peptide, conserved in all six RLD2-motif structures solved in this work. **B.** Detail of the RLD2-DxDKDxD complex, showing contacts of the N-terminal part of the motif, involving the two buried water molecules. This view and the two next ones were prepared from the RLD2/BAZ2B structure (PDB :7Q42), which displayed the best resolution (1.95 Å). The binding mechanism is the same for all crystalized complexes. **C.** Detail of contacts involving the central lysine of the motif. **D.** Detail of contacts involving the C-terminal part of the motif. Note the importance of ionic and polar contacts. Two water molecules buried at the interface mediate a number of critical H-bonds between several RLD2 residues and aspartates D1 and D3 of the motif. Hydrophobic contacts are also observed, with Tyr 3234 and Leu 3267 facing the hydrophobic side-chain of lysine K4 of the motif (C). Note that the two “x” amino acids are exposed to solvent and do not contribute to the interaction (B and D).

### The HERC2-UBE3A interaction is conserved across all vertebrates and most animals with a CNS

HERC2 is found in most animal species, yet not in fungi or more distant organisms. By aligning RLD2 HERC2 domains across a diverse group of organisms, we found that the DxDKDxD-binding surface of HERC2 is highly conserved across most animal species (Figure 5A). UBE3A proteins are found in most animals as well as in several fungi (Figure 5B). However, we found that UBE3A proteins display the HERC2-binding DxDKDxD motif only in animals that possess a brain or CNS, such as vertebrates, molluscs, or arthropods (Figure 5B). This correlation further suggests that the HERC2-UBE3A interaction may be involved in the development and/or function of the animal brain. By native holdup, we found that the human HERC2 RLD2 domain bound with a similar affinity (K_d_ ∼ 0.1 μM) to endogenous UBE3A from human (Figure 2B), mouse (Figure 5C and supplementary Figure 4) and zebrafish (Figure 5D and supplementary Figure 5). This indicates that not only this interaction, but also its binding strength, are under strong evolutionary pressure across vertebrate species.

**Figure 5.**
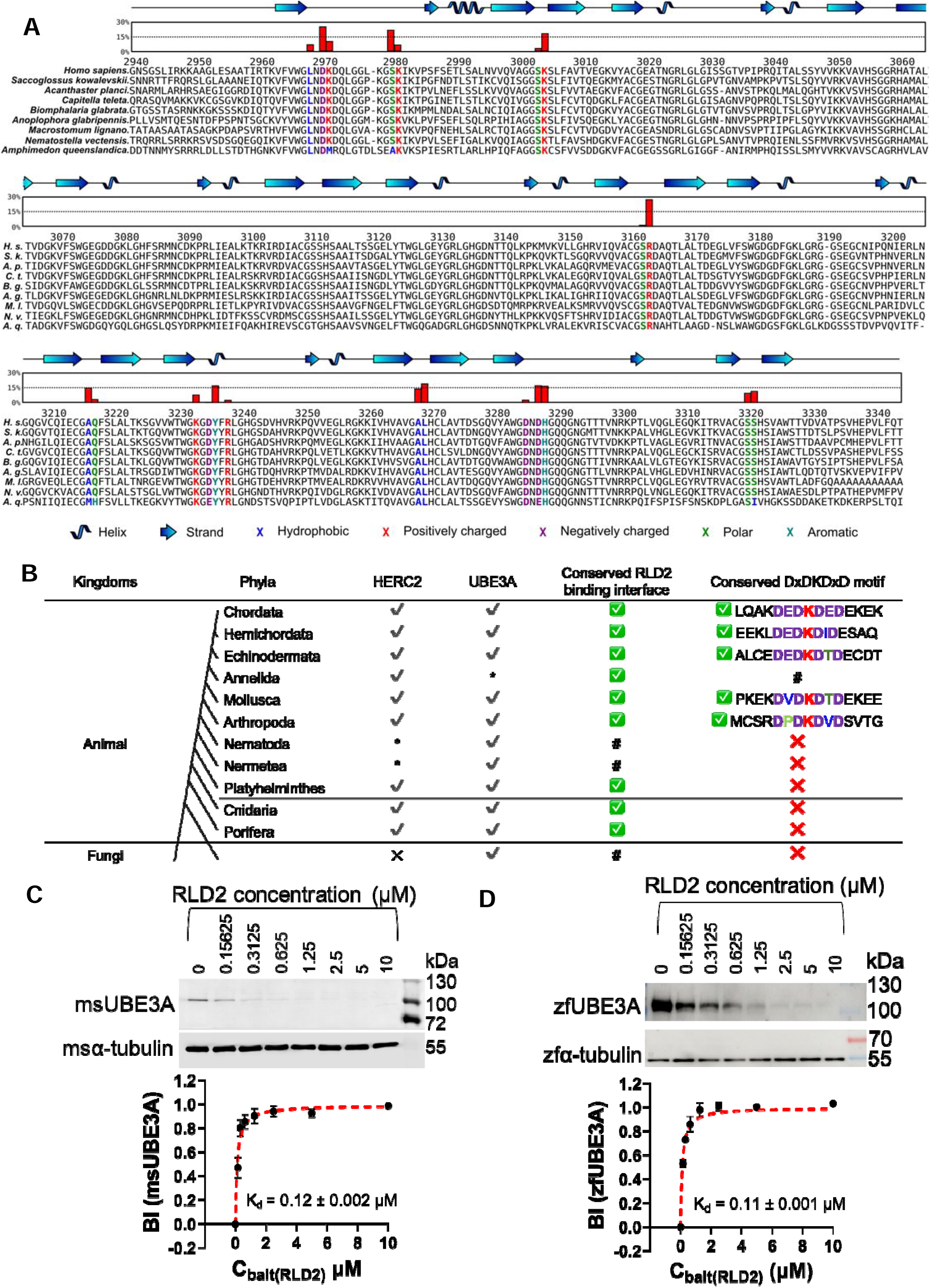
Conservation of HERC2 RLD2 domain across different species and of DxDKDxD motifs across different species and proteins. **A.** Sequence alignment of HERC2 RLD2 domain of species from 9 different animal phyla. Blades are structurally aligned based on their secondary structure elements, displayed on top of the sequences. The parts of the beta strands oriented towards the “top” side of the propeller supporting the bound peptide motif are colored in dark blue. For each amino acid of the domain, the percentage of accessible surface area buried by the bound peptide is plotted on top of the sequence alignment (red bars). Amino acids buried by the peptide are colored according to physicochemical properties. RLD2 amino acids are numbered according to the human HERC2 UniProt sequence. **B.** Conservation of HERC2 RLD2 domain and UBE3A DxDKDxD motifs across proteomes of different phyla. ?D: proteins are conserved across phyla; ?: proteins are not conserved across phyla; *: proteins did not appear in the search; ?: interface between HERC2 and UBE3A are conserved; ?: interface between HERC2 and UBE3A are not conserved; #: unknown; ---: separator between animal with and without central nervous system. **C-D.** Native holdup between human RLD2 and UBE3A from mouse and zebrafish. Biotinylated MBP-RLD2 was titrated from 10 μM to 0 μM to capture endogenous full-length UBE3A protein from lysates of mouse brains (P8) (**C**) and zebrafish embryos (2dpf) (**D**). Please see Supplementary Figures 4 & 5 for full blots of holdup titrations.

### HERC2 binds to a handful of other DxDKDxD-containing proteins

Phylogenetic sequence analysis also revealed that HERC2 proteins retain a DxDKDxD-binding surface in animals devoid of a CNS, where UBE3A no longer displays a DxDKDxD motif (Figure 5B). This suggests that the RLD2 domain of HERC2 may bind to other candidate DxDKDxD motifs found within proteins distinct from UBE3A. A sequence search of the entire human proteome using SlimSearch identified only 14 proteins containing the perfect DxDKDxD motifs (Supplementary Table 3). Nine of these proteins, including UBE3A, are involved in neuronal and/or neurodevelopmental syndromes (Figure 6A). DOCK10, in particular, is a Guanidine Exchange Factor (GEF) belonging to the DOCK-D family of RhoGEFs that is highly expressed in the brain [18]. It activates the small Rho GTPases RAC1 and CDC42, thereby promoting cytoskeletal remodeling and regulating dendritic spine formation in early post-natal neurodevelopment [14]. The deletion or mutation of DOCK10 has been documented in some cases of ASD [19]. PCM1 (Pericentriolar Material 1) and USP35 are cell cycle regulators and their dysregulation can disrupt neurodevelopmental processes [20–23]. ARID4A, RERE, BAZ2B and ARIP4 are chromatin remodelers and their dysregulations are related to neurodevelopmental disorders [24–34]. In all the aforementioned proteins, the DxDKDxD motif is conserved across most vertebrate species and is located in a predicted disordered region, which favors its availability for partner binding.

**Figure 6:**
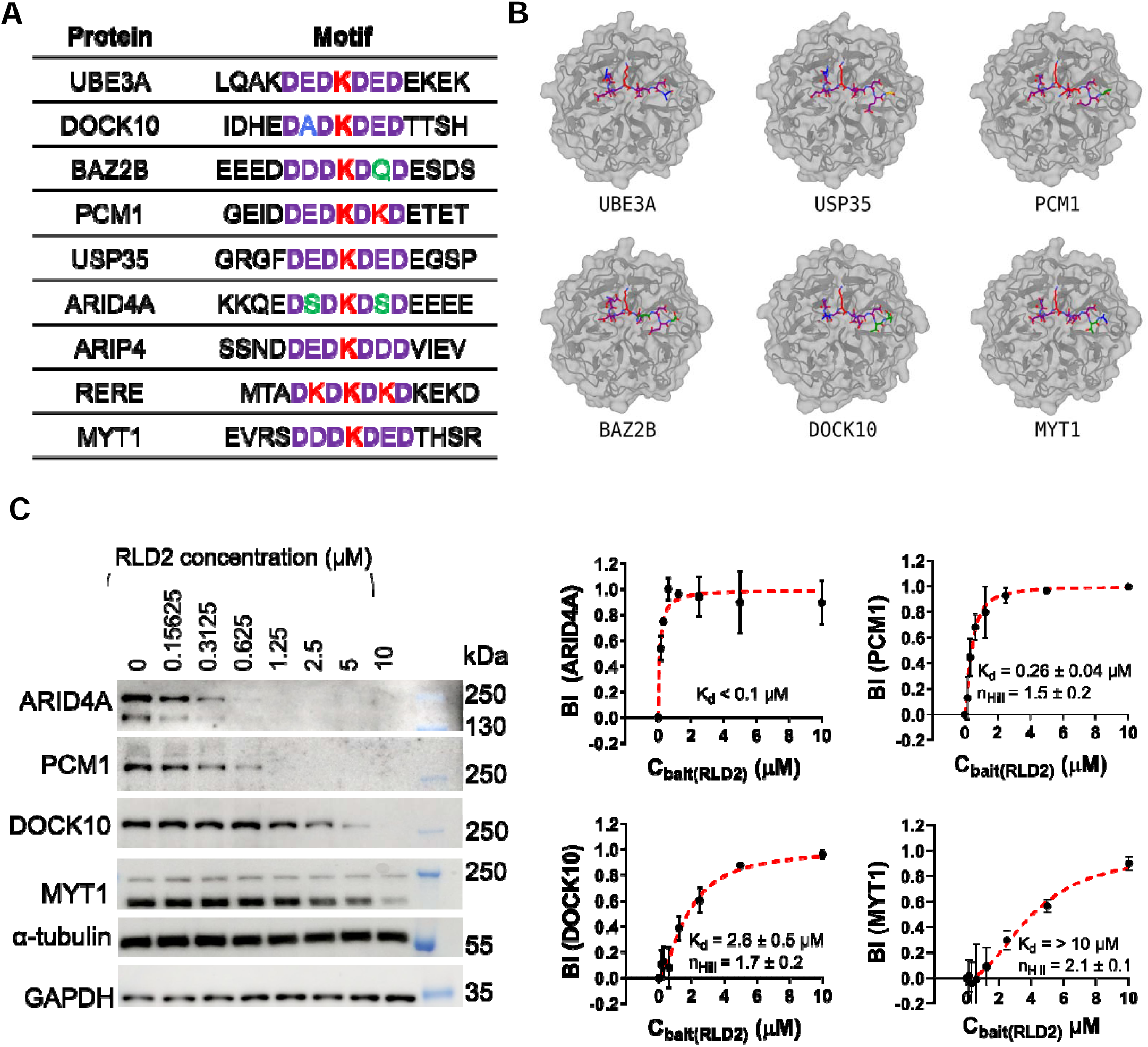
HERC2 binds several DxDKDxD motif containing proteins. **A.** Presence of DxDKDxD motif in other human proteins. **B.** Structures of complexes between HERC2 RLD2 domain and the DxDKDxD motifs of six different human proteins. Please see Supplementary Table 2 for the data collection and refinement parameters of the crystal structures. **C.** Native holdup titrations. The purified recombinant RLD2 domain of HERC2 was immobilized on streptavidin resin and the depletion ratio of different proteins was determined from lysates of SH-SY5Y cells. Binding curves derived from the holdup titrations, fitted based on a hyperbolic binding equation or Hill-equation based on better fit (red dashed line) to determine the binding affinity. K_d_: equilibrium binding dissociation constant. Please see Supplementary Figures 7 and 8 for full blots of native holdup titrations.

Therefore, we moved on to solve the structures of RLD2 bound to five of the aforementioned DxDKDxD motifs, derived from DOCK10, PCM1, USP35, BAZ2B and MYT1 (Figure 6B). The presence of side chains of variable bulkiness at the non-conserved “x” positions, and the higher resolution of some of the structures (culminating at 1.95 Å for the RLD2-BAZ2B complex) allowed us to ascertain the orientation and position of the peptide in the pocket, which was strictly conserved in all the RLD2-DxDKDxD complexes, including the initial RLD2-UBE3A structure.

We then used the competitive fluorescence polarization assay to measure the affinity of the HERC2 RLD2 domain for DxDKDxD motif-containing peptides derived from DOCK10, PCM1, USP35, BAZ2B, MYT1, ARID4A, ARIP4 and RERE. Most peptides bound in a comparable affinity range (1 µM < K_d_ < 30 µM) (Supplementary Figure 6 and Supplementary Table 4). We also quantified the affinities of the HERC2 RLD2 domain to endogenous full-length ARID4A, PCM1, DOCK10 and MYT1 by native holdup assay (Figure 6C, Supplementary Figure 7 & 8). Remarkably, competitive FP for RLD2 binding to DxDKDxD peptides and native holdup for RLD2 binding to endogenous full-length proteins showed similar affinities of RLD2 for DOCK10 (K_d (cFP)_ = 2.96 ± 0.17 µM) µM vs K_d (holdup)_ = 2.60 ± 0.5 µM) and MYT1 (K_d (cFP)_ = 25.46 ± 5.61 µM vs K_d (holdup)_ > 10 µM) but showed quite different affinities for PCM1 (K_d (cFP)_ = 6.93 ± 0.23 µM vs K_d (holdup)_ = 0.26 ± 0.04 µM) and ARID4A (26.58 ± 2.23 µM vs K_d (holdup)_ < 0.1 µM) (compare Figure 6D and Supplementary Table 4). This suggests that both DOCK10 and MYT1 essentially bind HERC2 RLD2 through their identified DxDKDxD motif. In contrast, PCM1 and ARID4A may contain additional regions that contribute to the binding to RLD2. Indeed, in ARID4A, we noticed the presence of a second region containing an “imperfect” DSEKDEK sequence which also displayed significant binding to HERC2 RLD2 in cFP assay (Supplementary Table 4). The combination of these two HERC2 binding sites in ARID4A might increase the binding affinity through avidity effects.

### Interaction of DOCK10 with HERC2 modulates DOCK10-mediated activation of RAC1 and CDC42, as well as its effect on dendritic spine morphogenesis

We further investigated the interaction of HERC2 with DOCK10, a RhoGEF known to regulate dendritic spine formation at critical stages of post-embryonic neurodevelopment [14]. We confirmed by co-immunoprecipitation experiments in HEK293T cells that the exogenous GFP-fused HERC2 RLD2 domain binds exogenous mCherry-fused full-length DOCK10 (Figure 7A and Supplementary Figure 9). Moreover, site-directed mutagenesis altering either the peptide binding groove of RLD2 or the DxDKDxD motif of DOCK10 disrupted this interaction (Figure 7A). Furthermore, mutagenesis of the DxDKDxD motif in the N-terminal region of DOCK10 led to significantly impaired RhoGEF activity of DOCK10 towards CDC42 and RAC1 (as measured by GTP-CDC42 and GTP-RAC1 pulldown assays) (Figure 7B and 7C and Supplementary Figure 10A-B). This was remarkable, given that the GEF activity of DOCK10 is located in its distal DHR2 domain, at the C-terminus of the protein. Next, we repeated these DOCK10-mediated GTPase activation assays in the absence of HERC2. HERC2-specific siRNA efficiently knocked down HERC2 expression and this was concomitant with a 40% decrease in DOCK10-mediated activation of CDC42 and RAC1 (Figure 7D and Supplementary Figure 10C-D). This suggested that HERC2 interaction with DOCK10 stimulates DOCK10 GEF activity. Notably, siRNA-mediated knockdown of HERC2 had no detectable impact on the GEF activity of TRIO, another RhoGEF known to activate RAC1 (Supplementary Figure 10E-F) [35]. Finally, mutagenesis of the DxDKDxD motif of DOCK10 also disrupted the ability of DOCK10 to promote dendritic spine formation in transfected cultured murine hippocampal neurons (Figure 8). These results suggest that HERC2 binding to the DxDKDxD motif of DOCK10 can positively stimulate DOCK10’s GEF activity and subsequent control of dendritic spine morphogenesis.

**Figure 7:**
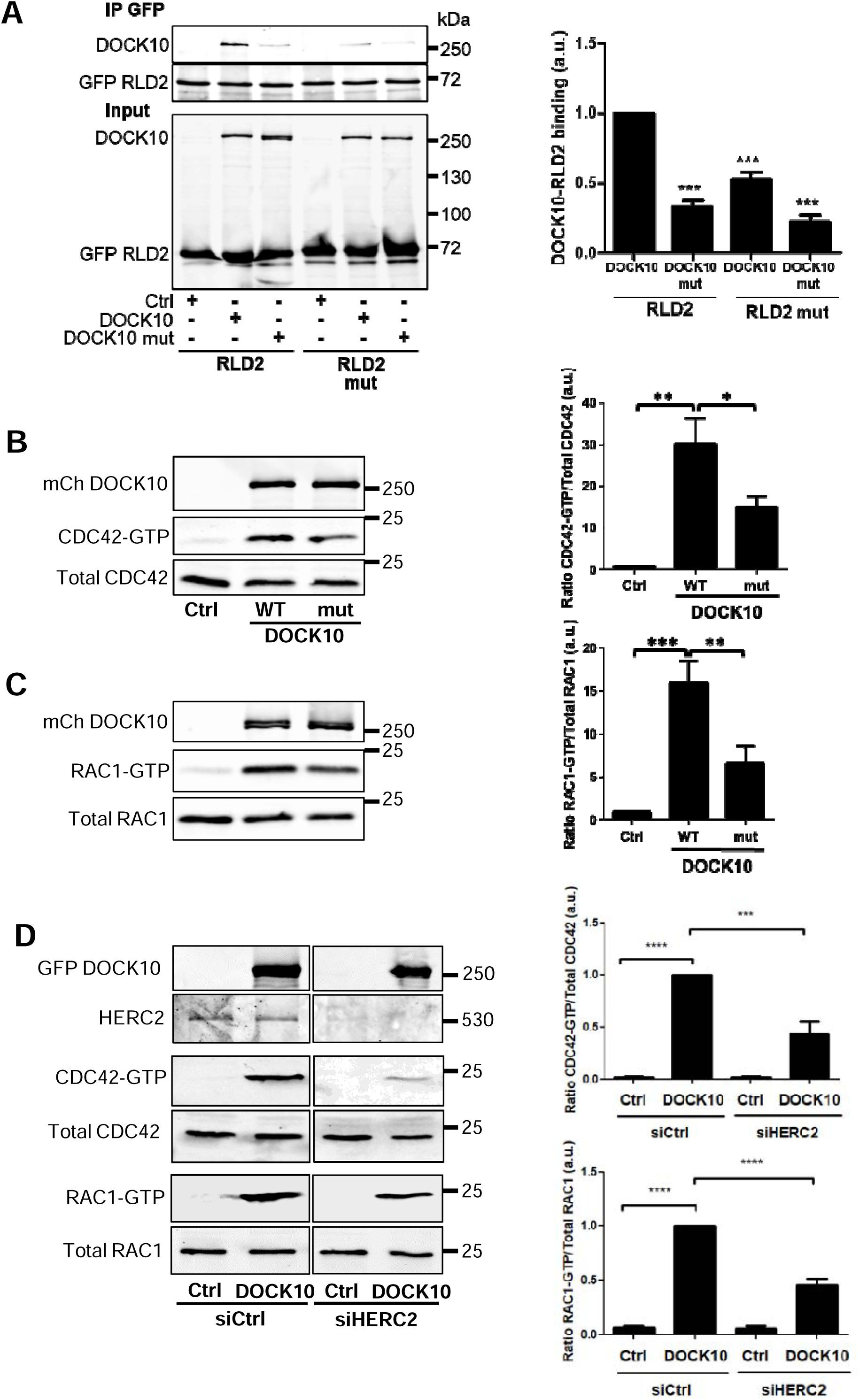
The interaction between DOCK10 and RLD2 is required for DOCK10-mediated activation of CDC42 and RAC1. **A.** Western Blot analysis of a co-immunoprecipitation assay of GFP-RLD2 and mCherry-DOCK10. HEK293T cells were transfected with GFP-RLD2 (or GFP-RLD2 mut, mutated on the DxDKDxD-binding interface as described in methods) and mCherry-DOCK10 (or mCherry-DOCK10mut, mutated on the DxDKDxD motif as described in methods) or a control empty vector. RLD2 was immunoprecipitated using an anti-GFP antibody, and the co-precipitating DOCK10 was detected using a mCherry antibody. **B.** Histogram of the quantification of the experiment presented in A. Data are presented as the mean ± SEM of 5 independent experiments. Statistical analyses were made using one-way ANOVA followed by Dunnett’s test. Asterisks indicate datasets significantly different to the wt interaction between RLD2 and DOCK10 (***p<0.001). Please see Supplementary Figure 9 for full blot. **B-D.** CDC42-GTP and RAC1-GTP pull-down assays were performed in HEK293T cell lysates (transfected as described below). Active GTP-CDC42 (or GTP-RAC1) was affinity purified using the CDC42/RAC1-interactive binding (CRIB) domain of PAK1, immobilized on Glutathione-Sepharose beads. Purified GTP-bound and total CDC42 (or RAC1) were detected by Western blot, using an anti-CDC42 antibody (or anti-RAC1 antibody). Protein expression in the cell lysates was verified by immunoblotting with an anti-mCherry antibody to detect DOCK10 and an anti-HERC2 antibody (D). One representative experiment (Western Blot) is shown on the left-hand panels. Right-hand panels: histogram of the quantification of at least 4 experiments. **B.** CDC42-GTP pull down assay performed on HEK cells transfected with mCherry DOCK10, mCherry DOCK10mut or a control vector. CDC42 activation mediated by wt DOCK10 was arbitrarily set to 100%, in order to be able to compare the individual experiments. The % of CDC42 activation was calculated from at least 4 independent experiments (mean ±SEM). Statistical analyses were made using the non-parametric two-tailed Mann-Whitney test (*p<0.05, **p<0.01, ***p<0.001). **C.** RAC1-GTP pull down assay. Experiments were performed and quantified as described in B, except that an anti-RAC1 antibody was used for immunoblotting. **D.** CDC42-GTP and RAC1-GTP pull down assays performed in the absence of HERC2. HEK293T cells were co-transfected with siHERC2 or siCtrl and pEGFP-DOCK10, pEGFP-DOCK10mut or a control vector (siRNA transfected for 72 h, cDNA transfected for 48 h). Active GTP-CDC42 (upper panels) or GTP-RAC1 (lower panels) were affinity purified as described in B. Purified GTP-bound and total CDC42 or RAC1 were detected by Western blot, using the relevant anti-CDC42 and anti-RAC1 antibodies. Protein expression of DOCK10 and HERC2 in the cell lysates was verified by immunoblotting with an anti-GFP and anti-HERC2 antibody. The % of CDC42 (or RAC1) activation was calculated from at least 4 independent experiments (mean ±SEM). Statistical analyses were made using the non-parametric two-tailed Mann-Whitney test (*p<0.05, **p<0.01, ***p<0.001, ****p<0.0001). Please see Supplementary Figure 10 for full blots.

**Figure 8:**
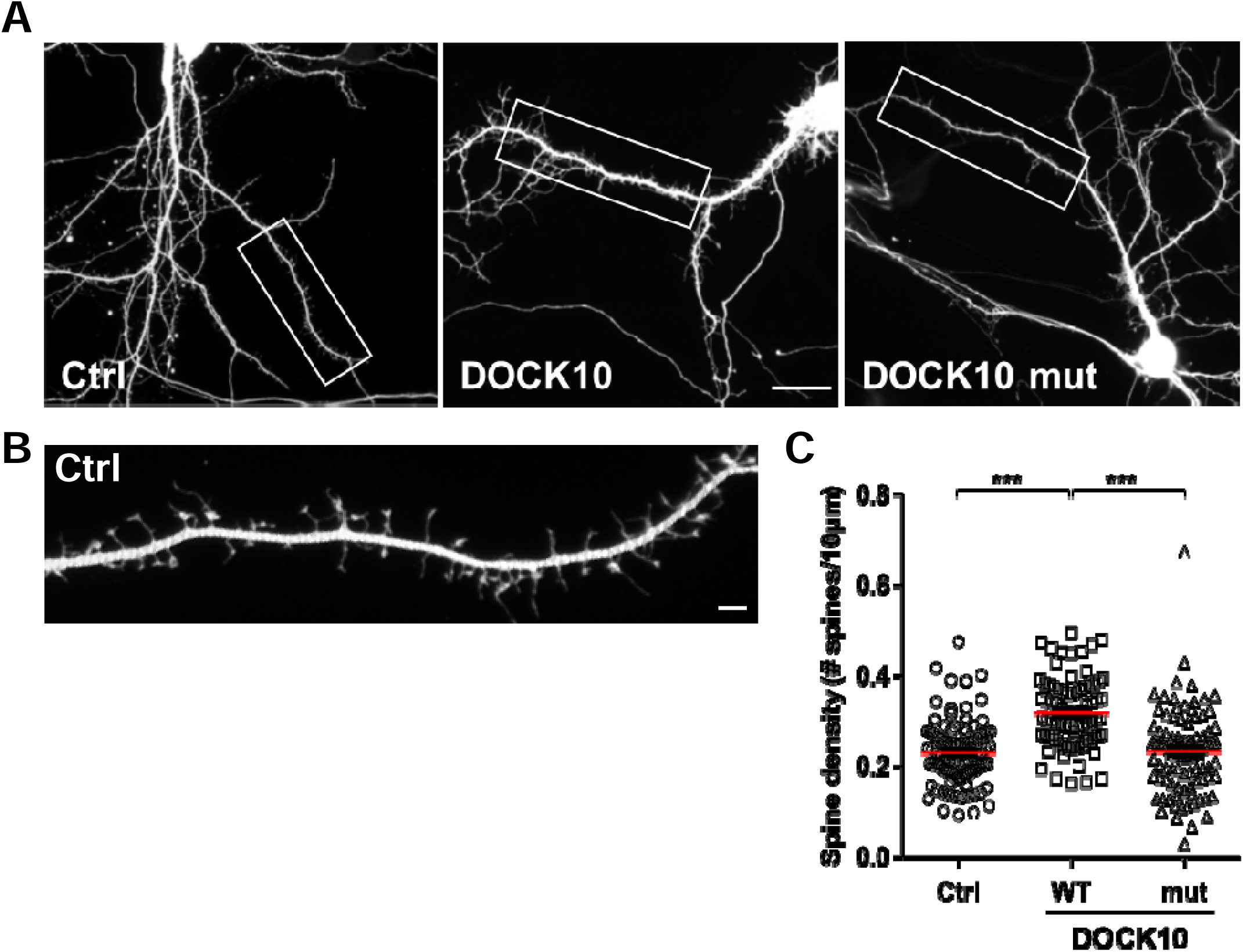
Expression of DOCK10mut, defective in RLD2 binding, leads to decreased dendritic spine formation in primary hippocampal neurons. **A.** Representative micrographs of cultured murine hippocampal neurons transfected with mCherry DOCK10, mCherry DOCK10mut or a control vector. Neurons were transfected at 8 DIV and fixed at 11 DIV (experimental conditions to reveal spine formation). Scale bar: 20 µm. **B.** Higher magnification images of the regions (white rectangles) selected in A, illustrating the dendritic spine morphology and density in the neurons transfected as described in A. Scale bar: 1 µm. **C.** Quantification of dendritic spine density of hippocampal neurons transfected as described in A, measured by the number of spines/10 μm. Error bars indicate SEM (*n* = 3 independent experiments, with at least 15 neurons/condition/experiment).

### Structural analysis of the impact of HERC2 binding on the RhoGEF activities of DOCK10

We performed AI-based AlphaFold predictions of apo-DOCK10, DOCK10 bound to its partner GTPases RAC1, and DOCK10 bound to the RLD2 domain of HERC2 (Figure 9). All predictions consistently revealed a main folded unit consisting of DHR1-C2 domain (cyan-green), Armadillo helical repeats (green to orange) and the DHR2 GEF catalytical domain (red), reminiscent of that observed in experimental structures of DOCK2 and DOCK5 (Supplementary Figure 11). A long N-terminal extension (dark blue to lighter blue) wraps around the surface of the main folded unit. This extension includes a kinked α-helix at the extreme N-terminus, an αβαβα structure contiguous to the DHR1-C2 domain and a PH domain contiguous to the DHR2 domain (Supplementary Figure 12). Apo-DOCK10 is consistently predicted in an extended, open conformation (Figure 9A). GTPase-bound DOCK10 systematically adopts a more compact, closed conformation (Figure 9B) where the RAC1 GTPase-bound DHR2 domain bends towards the armadillo core region. Notably, the predictions using either Rac1 or CDC42 gave essentially the same results. Finally, RLD2-bound DOCK10 adopts a more fuzzy, intermediate open-closed conformation, where the DHR2 domain explores various positions relative to the rest of the structure (Figure 9C). RLD2 recognizes DOCK10 via its DxDKDxD motif, supplemented by a large additional interface.

**Figure 9:**
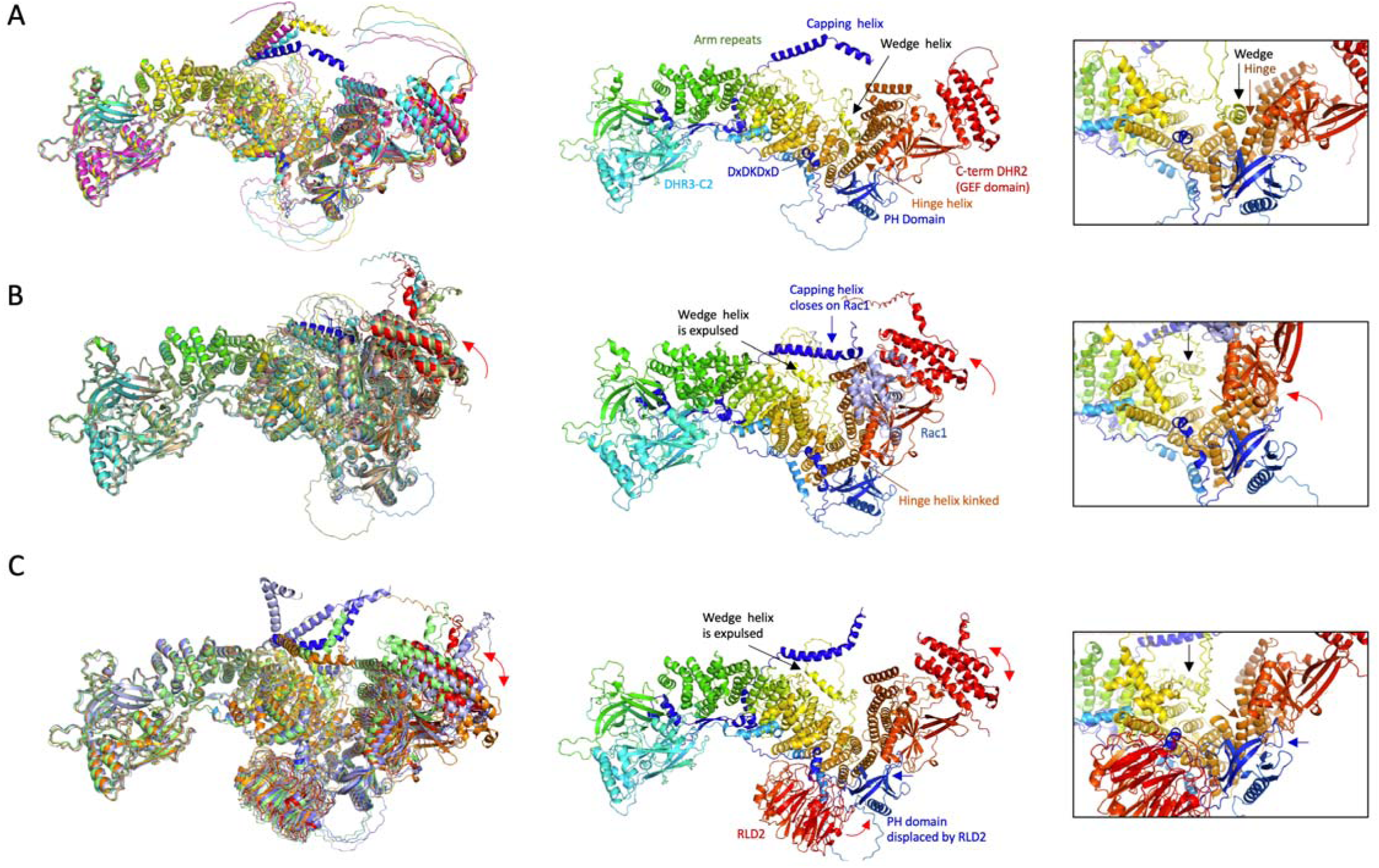
Structure predictions provide a structural basis for the activation of the GEF activity of DOCK10 by HERC2. **A.** Modeled structure of full-length apo-DOCK10. Left: the 5 superimposed structures of full-length human DOCK10, modeled by AlphaFold 3 predictor. All structural features, including the long N-terminal extension that wraps around the main folded body, are predicted with high reproducibility. Middle: the best ranked structure, colored blue to red from N-term to C-term. Structural elements discussed in the main text are indicated. Right: zoomed view of the wedge helix (yellow) inserted within the last repeat of the armadillo body and the long straight hinge helix (orange). **B.** Modeled structure of Rac1-bound DOCK10. Left: the 5 superimposed structures of Rac1-bound DOCK10, modeled by AlphaFold 3. Middle: the best ranked structure. Rac1 is colored pale blue. The DHR2 GEF catalytic domain undergoes a whole body displacement towards the rest of the structure. The N-terminal helix closes upon the bound Rac1. Right: zoomed view showing the wedge helix (yellow) expulsed from the armadillo body and the hinge helix undergoing a kink, both supporting the whole-body movement of DHR2 and subsequent capping by the N-terminal helix of DOCK10. **C.** Modeled structure of RLD2-bound DOCK10. Left: the 5 superimposed structures of RLD2-bound DOCK10, modeled by AlphaFold 3. All structural features, including the pose of the RLD2 domain, are predicted with high reproducibility. However, the DHR2 catalytic domain now explores a variety of intermediate poses relative to the rest of the protein. Middle: the best ranked predicted structure. HERC2 RLD2 domain (red) binds to the DxDKDxD motif (dark blue) and also established additional interface with other regions of DOCK10, including the PH domain, which is slightly displaced in direction of the armadillo body. Right: The wedge helix (yellow) is expulsed in the best prediction and in two other ones, or distorted in the two remaining ones. The hinge helix remains mostly straight with however a slight bend in some of the predictions. These predictions support a model where RLD2 binding induces a push upon the PH domain, which in turn favors the “closing” movement of DHR2 towards the rest of the structure, including expulsion of the wedge helix. This conformational change favors the recruitment of Rac1, thereby stimulating the GEF activity.

Three outstanding features appear to account for the observed conformational differences. (i) GTPase capping motif: in both the apo-DOCK10 and the RLD2-bound structures, a helical stretch located at the extreme N-terminus of DOCK10 stands free in the solvent, away from the main core structure of DOCK10 (“Capping helix” in Figure 9A and 9C). In the GTPase-bound structure, this helical stretch systematically “caps” the bound GTPase in the vicinity of switch 2, a conformationally labile loop relevant for Mg^2+^-dependent binding of Rho-GTPases to nucleotides (Figure 9B) [36, 37]. The critical residues of this GTPase-capping motif are strongly conserved in DOCK10 and the other two members of the DOCK-D subfamily of RhoGEFS, DOCK9 and 11 (Supplementary Figure 13). (ii) Wedge helix: in apo-DOCK10, a long-disordered loop connecting two armadillo repeats includes a short helical stretch (sequence K1264DVLNSIAAFS1274) that acts like a wedge that inserts between two helices of another Arm repeat (Figure 9A). In GTPase-bound DOCK10 structures, this wedge helix is systematically expulsed (Figure 9B). This allows the two surrounding helices to come in closer contact to each other, contributing to the closing of the structure. In the RLD2-bound DOCK10 structures, the wedge helix is either expulsed or only partly inserted (Figure 9C). (iii) Hinge helix: in apo-DOCK10, a long straight helix connects the end of the solenoid structure to the A lobe [36, 37] of the DHR2 domain. In GTPase-bound DOCK10, this helix undergoes a kink at position T1668, thereby promoting the displacement of the entire DHR2 domain (Figure 9B). In the RLD2-bound structures, this helix remains straight (Figure 9C), indicating that its kinking and the subsequent full compaction of DOCK10 become effective only when the DHR2-bound GTPase needs to reach for the N-terminal capping motif.

Altogether, these observations support a model whereby the HERC2 RLD2 domain activates the GEF activity of DOCK10 by promoting its binding to GDP-bound GTPases. In this model, RLD2 binds to DOCK10 *via* its DxDKDxD motif plus a large additional interface. This induces conformational changes that favor the expulsion of the wedge helix, resulting in higher mobility of the DHR2 domain. The latter is then allowed to explore a position compatible with high-affinity charging of the GDP-bound GTPase, which involve interface contacts with both the C-terminal DHR2 domain and the N-terminal capping motif of DOCK10. Once charged, the GDP-GTP exchange and subsequent discharge of the GTP-bound, activated GTPase, can proceed.

## Discussion

*UBE3A* and *HERC2* genes are altered in neurodevelopmental syndromes presenting overlapping clinical features, and the UBE3A and HERC2 proteins interact with each other. This prompted us to investigate the molecular details of the HERC2-UBE3A complex. We showed that the HERC2-UBE3A complex boils down to a domain-motif interaction involving the RLD2 domain of HERC2 and a short DxDKDxD motif of UBE3A. We present evidence that this complex is present in most animals with a CNS. Furthermore, we identified and validated a handful of additional HERC2-binding proteins that contain a *bona fide* RLD2-binding DxDKDxD motif. The ensemble of our observations thus points to a network of proteins relevant for brain development that share the ability to bind to a defined domain of the giant protein HERC2.

In this work, we have devoted particular effort not only to the demonstration and structural characterization of protein-protein interactions but also to the determination of their binding dissociation constants (K_d_), which are essential quantitative parameters for the accurate description and biological interpretation of protein networks. For this purpose, we have mainly used the native holdup assay [16], which allows the precise determination of binding affinities of any chosen molecule saturated on resin, against any full-length endogenous protein detectable by Western blotting within a biological cell extract. This alleviates many of the problems associated with most traditional *in vitro* approaches to affinity determination, which require the use of pure and fully active proteins. In particular, this approach allowed us to measure binding constants between UBE3A and full-length HERC2, an extremely large protein that is difficult to purify in a native form. The holdup assay also allowed us to demonstrate the strong conservation of the HERC2-UBE3A affinity constant across distant vertebrate species. Finally, using a combination of holdup and fluorescence polarization assays, we were able to distinguish cases in which the binding affinities of entire proteins and of their minimal interacting fragments are similar (e.g. HERC2-UBE3A and HERC2-DOCK10 complexes), from cases in which the binding affinities of the fragments are strongly diminished as compared to the entire proteins. These observations suggest that distal regions or multimerization effects may additionally contribute to complex formation (e.g. HERC2-PCM1 and HERC2-ARID4A complexes).

Remarkably, HERC2 and nine of its DxDKDxD-containing partners have documented implications in normal and/or pathological neurodevelopment through various cellular functions. Four of them are chromatin remodelers (ARID4A, BAZ2B, RERE, and ARIP4). This is striking because HERC2 itself is a strong candidate for chromatin remodeling as it contains a ZZ domain that specifically interacts with Histone H3 [38]. In addition, *HERC2* mutations were recognized early on to induce sperm defects in mice [39]. The DNA-binding transcriptional regulator ARID4A is known to directly regulate the epigenetic modification of the AS/PWS IC (Angelman Syndrome / Prader Willi Syndrome imprinting center) that controls the expression of *UBE3A*-encoding alleles [40]. Interestingly, we found that full-length ARID4A binds the RLD2 domain of HERC2 with a strong affinity comparable to that of UBE3A. Among the other DxDKDxD-containing chromatin remodelers, the haploinsufficiency of *BAZ2B* is strongly associated with developmental delay, intellectual disability, and autism spectrum disorder [33, 34, 41]. ARIP4/RAD54L2 is a steroid hormone receptor co-factor involved in chromatin remodeling and Androgen Receptor regulation in sex differentiation and spermatogenesis [42]. It is also a major binder of the DYRK1A kinase, which is implicated in neuronal differentiation defects in ASD and Down Syndrome [32, 43, 44]. MYT1 is a cell cycle kinase [45] and proneuronal transcription factor [46] that binds to the CoREST protein [44] to serve as a neuron-specific subunit of the LSD1/CoREST chromatin remodeling complex [47, 48]. Deletion of *MYT1* gene is also associated with intellectual disability [27].

Other DxDKDxD-containing proteins include RERE, a transcriptional co-repressor that modulates Histone deacetylases 1/2 and is important for proper patterning of gene expression during early development [49]. Mutation of RERE is associated with the NEDBEH neurodevelopment disorder [29]. The DxDKDxD-containing protein USP35 is a de-ubiquitinase involved in various cancers [23, 50–53] and has recently been suggested to play an active role in the development of ciliopathies [54].

PCM1 was ranked among the top RLD2-binders in our affinity measurements. PCM1 was previously shown to interact with HERC2 [10, 55], and the binding site to PCM1 was restricted to a HERC2 fragment encompassing the RLD2 domain [10], confirming that the DxDKDxD motif is a hallmark for HERC2 RLD2-binding. PCM1 has been proposed to be involved in microtubule organization during early neuronal development [56]. Mutations of PCM1 have been associated with schizophrenia [57], and PCM1 KO mice exhibit progressive ciliary, anatomical, psychomotor, and cognitive abnormalities [57, 58]. The PCM1-HERC2 interaction is also remarkable in that HERC2 and UBE3A have both been reported to localize to the centrosome at certain steps of the cell cycle [59, 60]. It is tempting to speculate that an altered dosage of HERC2/PCM1 complexes might alter some of the potential functions of PCM1 in neurodevelopment.

Finally, we were particularly intrigued by the DOCK10 RhoGEF, which contains a DxDKDxD motif within a small, presumably disordered stretch of its N-terminal region, located distal to the C-terminal DHR2 domain responsible for GEF activity towards CDC42 and RAC1. DOCK10 has previously been implicated in dendritic spine formation during the early steps of postnatal mouse brain development [14] but how its GEF activity is regulated remains unknown. The GEF activity of DOCK10 was impaired upon depletion of HERC2, and mutating the DxDKDxD motif in DOCK10 strongly affected both its GEF activity and its ability to promote dendritic spine formation. This study therefore identifies HERC2 as the first regulator of DOCK10 activity in neurons.

Structural predictions provided us with a convincing model of activation of DOCK10 GEF activity upon binding to HERC2, in full agreement with our experimental data. In contrast to the well-described regulation of DOCK-A and -B subfamily GEFs by binding to the adapter protein ELMO, a regulatory mechanism for DOCK-D subfamily GEFs has not been described so far. ELMO binds to the N-terminal SH3 domains of DOCKA/B, bringing its PH domain in close proximity to the catalytic DHR2 domain of the GEF. This leads to release of the autoinhibitory status of DOCKs and exposure of their DHR2 domain for RAC activation. DOCK-D members, in contrast, lack this N-terminal SH3 domain and therefore do not bind the ELMO adapter. Uniquely, the DOCK-D proteins, including DOCK10, contain a PH domain instead at their N-terminus, which is spatially proximal to the DHR2 domain of DOCK10, and could function similarly to the PH domain of ELMO. Our model proposes that the RLD2 domain of HERC2, by binding to the DxDKDxD motif of DOCK10, induces a push on the nearby PH domain, which in turn induces a conformational change favoring the recruitment of RAC1, and hence its activation. Our model therefore proposes the first molecular mechanism by which DOCK10 might be activated.

More generally, our data suggest that HERC2 forms with all its DxDKDxD partners a tightly balanced network, where the relative abundance of each complex is relevant to optimal brain development. Under homeostatic conditions, HERC2 can be expected to form complexes in varying proportions with the different DxDKDxD partners discussed above, depending on the tissue and cell type, the relative amounts of HERC2 and each partner, their localization and their conformational states, which may be regulated by third-party binders or post-translational modifications. The strong conservation of the affinity constant observed here for binding of HERC2 to UBE3A from fish to mammals suggests that the proportions of the different complexes involving the HERC2 RLD2 domain have been finely tuned and conserved during evolution. Under pathological conditions, where one member of the network (HERC2, UBE3A or another DxDKDxD motif protein) is missing or defective due to genetic alterations, an imbalanced network may arise, in which complexes will be redistributed in anomalous proportions. For instance, this might lead to a change in the abundance of HERC2-DOCK10 complexes, which in turn might lead to altered dendritic spine formation. In Angelman Syndrome brain tissues, where UBE3A is missing or misfolded, HERC2 is likely to form more abundant complexes with its other potential partners (e.g., HERC2-DOCK10 complexes), thereby altering their conformation, localization, availability for other interactions, and so forth. In brain tissues where UBE3A is expressed at higher amounts (Dup15q Syndrome), an abnormal redistribution of the network is also expected. Subsequently, the activities of DxDKDxD proteins may be affected in brains with altered UBE3A expression. Hence, we suggest that proper dosage of HERC2-DxDKDxD complexes and conformational changes that DxDKDxD proteins may undergo upon binding to HERC2 might be important for proper brain neurodevelopment.

## Material and methods

### Sequence analysis

To identify the different orthologs, several strategies were employed. First, using the NCBI tool, we searched for similar proteins based on global architecture [61]. This approach was effective for UBE3A, as it is the only protein that contains the Azul and HECT domains. However, this strategy may overlook sequences that are poorly annotated. Therefore, we performed a systematic BlastP search [62] against protein databases from various phyla individually.

To map the sequence conservation of the RLD2 domain structure across different species, we used the conservation tool in ChimeraX [63], which performs multiple sequence alignments (MSA) using Clustal Omega [64] to calculate the AL2CO score [65]. Searches for the motif (DxDKDxD) were performed using SlimSearch software [66] (accessed in 2019). This software identifies proteins containing the motif in unfolded regions, which are favorable for motif-domain interactions.

### Accessible surface area and accessibility

The “Accessible Surface Area and Accessibility Calculation for Protein” software (version 1.2) (Available at: http://cib.cf.ocha.ac.jp/bitool/ASA/ (Accessed in 2020)) was used to calculate the percentage of the surface area of each amino acid in the RLD2 domain that was buried by the DxDKDxD-containing peptides. To do this, the solvent-exposed surface area of each amino acid of the RLD2 domain was compared in the presence and absence of the peptide. Since the structure with the BAZ2B peptide provided the highest resolution, it was selected for this comparison. This comparison enabled the identification of the regions of the RLD2 domain that were masked by the peptide upon interaction.

### Peptide selection and production

Peptide sequences were defined after the SlimSearch screen mentioned above. Peptides for crystallization and fluorescence polarization measurements were synthesized and purified by Pascal Eberling at the peptide synthesis platform of IGBMC. Peptide sequences can be found in Supplementary Tables 1 and 3.

### Constructs

The constructs used for crystallization and fluorescence polarization assays were cloned into the pETM41 vector, which carries a His_6_-MBP-TEV tag fused to the N-terminus of the HERC2 RLD2 domain (Uniprot: O95714, residues 2941-3342) and full-length UBE3A (Uniprot: Q05086-2).

For native holdup assays, constructs were cloned into the same pETM41 vector but with an AviTag-His_6_-MBP-TEV fusion. The AviTag enables *in vivo* biotinylation, while His_6_-MBP facilitates purification via affinity chromatography and enhances solubility. The TEV cleavage site allows tag removal, yielding untagged recombinant proteins for crystallization and fluorescence polarization assays.

### Protein expression and purification

All constructs were expressed in *Escherichia coli (E. coli) BL21 (DE3)* competent cells. Transformed bacteria were selected with kanamycin. For biotinylated protein production, the AviTag constructs were co-transformed with *E. coli* biotin ligase (BirA) construct and selected with kanamycin and ampicillin. All expression cultures were performed with 0.2% glucose and shaking at 190 rpm (an excess amount of biotin (50 µM) was added for biotinylated protein production) and allowed to grow to OD600 of 0.5 at 37 °C before the temperature was reduced to 16°C. The expression was induced (OD600 = ∼1.0) by the addition to a final concentration of 1 mM IPTG (100 µM ZnSO_4_ was additionally added for UBE3A full-length) and left to grow overnight at 16°C. The expression cultures were harvested at 6000 xg, for 20 minutes at 4 °C. The biomass was frozen at -20 °C until the next step.

All biomasses were resuspended in lysis buffer containing 50 mM Tris-HCl pH 8.0, 400 mM NaCl, 5 mM glycerol, 5 mM beta-mercaptoethanol (β-ME), 1x cOmplete EDTA-free protease inhibitor cocktail, lysozyme (4 mg per 10 g wet biomass), 0.25 µg/ml DNase, and 0.25 µg/ml RNase. Cell lysis was performed by sonication for 1 minute 45 seconds (s) in 2 s on /2 s off cycles, 40 ml each time. Then, the lysate was centrifuged at 125 000 xg for 1 hour to obtain the clear lysate.

For RLD2, the proteins were subjected to nickel (His_6_-MBP-TEV construct) or amylose (AviTag-His_6_-MBP-TEV construct) affinity chromatography by loading the cleared lysate on a nickel column (HisTrap FF crude) or amylose resin (E8022, NEB) equilibrated with 20 mM HEPES pH 8.0, 300 mM NaCl, 3 mM β-ME. The unbound and non-specific bound contaminants were washed with the same buffer to which 40 mM imidazole was added to the washing step in the nickel column. The proteins loaded on nickel column or amylose resin were eluted by a buffer containing 250 mM imidazole or 10 mM maltose monohydrate, respectively. The eluted fractions were then subjected to SP Sepharose ion exchange chromatography. The proteins were eluted with NaCl gradient (from 100 mM to 1 M NaCl), containing 20 mM HEPES pH 8.0 and 3 mM β-ME. For native holdup assay, the proteins were polished with a size exclusion chromatography step (SEC) (HiLoad® 26/600 Superdex® 200 pg, Cytiva) in a buffer containing 20 mM HEPES pH 8.0, 300 mM NaCl and 3 mM β-ME. For crystallography, the MBP fusion was cleaved by TEV (“Tobacco Etch Virus”) protease and further purified by ion exchange chromatography (HiTrap™ SP HP, Cytiva) before polishing with SEC (HiLoad® 16/600 Superdex® 75 pg, Cytiva) in a buffer containing 20 mM HEPES pH 8.0, 300 mM NaCl and 3 mM β-ME. Protein was concentrated using centricon (10 kDa cutoff) to reach the concentration of 4 mg/mL.

For MBP and MBP-UBE3A full-length, the cleared lysates were applied on amylose resin equilibrated with buffer A containing 50 mM Tris-HCl pH 8, 300 mM NaCl, 3 mM β-ME. The unbound or non-specific bound fraction was washed with 2-column volume (CV) buffer A followed by 1.5 CV of buffer B containing 50 mM Tris-HCl pH 8, 100 mM NaCl, 3 mM β-ME. The proteins were eluted with buffer B containing 10 mM maltose monohydrate. MBP was applied directly on SEC column HiLoad® 26/600 Superdex® 200 pg (Cytiva), while MBP-UBE3A full-length was applied immediately on HiTrap Q HP (Cytiva) equilibrated with Buffer B and was subjected to anion exchange (performed at 2.5 ml/min). The unbound or non-specific bound contaminants were washed with wash buffer containing 50 mM Tris-HCl pH 8, 100 mM NaCl, 3 mM β-ME followed by 10 CV of gradient elution with buffer B containing NaCl concentration of 100 mM to 1 M NaCl. The fractions containing MBP-UBE3A full-length (∼45% gradient peak) was subjected to SEC (HiLoad® 26/600 Superdex® 200 pg, Cytiva). Monomeric MBP and MBP-UBE3A full-length were concentrated to 30 – 40 µM using centricon (10 kDa cutoff), snap-frozen with liquid nitrogen and stored at -80 °C for native holdup assays.

For UBE3A used in fluorescence polarization, proteins were first purified using affinity chromatography. The lysate was loaded onto a nickel column (HisTrap FF crude, Cytiva) in a buffer containing 50 mM Tris-HCl pH 8, 400 mM NaCl, and 3 mM β-ME, washed with 50 mM Tris-HCl pH 8, 100 mM NaCl, 35 mM imidazole, 3 mM β-ME, and eluted with the same buffer containing 80 mM imidazole. UBE3A was further purified using HiTrap Q HP (Cytiva) followed by SEC (HiLoad® 26/600 Superdex® 200 pg, Cytiva) as described above. Monomers were selected for MBP removal via overnight TEV protease cleavage at room temperature. Cleaved MBP (His_6_-tagged) was retained using nickel affinity chromatography (HisTrap FF crude, Cytiva) with a buffer containing 35 mM imidazole. A second ion exchange chromatography step (HiTrap Q HP) was performed as described previously. Finally, a SEC (HiLoad 16/600 Superdex 200 pg, Cytiva) in 20 mM Tris-HCl pH 8, 200 mM NaCl, and 3 mM β-ME yielded the purified monomeric protein. The protein was concentrated using a 10 kDa cutoff Centricon.

### Crystallization and structure determination

RLD2 (4 mg/mL) was used for crystallization assays. Drops were prepared using the sitting drop vapor-diffusion method in 96-well plates (200 nL protein solution + 200 nL reservoir) with a SPT Labtech Mosquito robot. For co-crystallization, RLD2 was incubated with the peptide at a 1:5 molar ratio for 30 minutes prior to crystallization. Both RLD2 apo crystals and RLD2 in complex with 6 different peptides (co-crystals) grew at 20 °C in a solution of tri-ammonium citrate 180 mM and 20% polyethylene glycol 3350.

To be tested for X-ray diffraction, crystals were transferred into a cryoprotectant solution (crystallization condition supplemented with 20% ethylene glycol) and subsequently flash-frozen in liquid nitrogen. Data collection was performed on the PROXIMA-1 and PROXIMA-2A beamlines at the SOLEIL synchrotron (Saint-Aubin, France) and on the X6DA beamline at the Swiss Light Source (Villigen, Switzerland).

Data were processed using XDS [67] (see supplemental data). The structure was solved by molecular replacement with Phaser (CCP4 software package) [68], using the third RLD domain of HERC2 (RLD3) structure (PDB: 3KCI, chain A) as the search model. Model building and improvement were carried out through iterative cycles of manual building in Coot [69] and refinements with Phenix [70]. The structure of RLD2 apo was deposited in PDB, with ID: 7Q40. The 6 structures of the complexes with peptides were deposited with PDB ID: 7Q41 (UBE3A), 7Q42 (BAZ2B), 7Q43 (DOCK10), 7Q44 (USP35), 7Q45 (MYT1) and 7Q46 (PCM1).

### Fluorescence polarization

All FP measurements were performed in a buffer containing 25 mM HEPES (pH 7.5), 150 mM NaCl, 1 mM Tris(2-carboxyethyl)phosphine (TCEP), and 0.005% Tween 20. Direct measurements were conducted by titrating RLD2 with a fixed amount of 50 nM fluorescent peptide (UBE3A 18-aa linked to fluorescein, Fluorescein-SLQAKDEDKDEDEKEKAA).

For the competitive measurements, the complex was formed by mixing 50 nM fluorescent peptide with 3.33 µM RLD2. Non-labeled peptides were then titrated against the fixed amount of the complex. Measurements were performed using a Pherastar plate reader. The different dilutions for direct and competitive measurements were transferred from 96-well plates to 384-well black plates in triplicate (10 µL per well). Fitting of the FP data was performed using Profit software [71].

### Cell culture

SH-SY5Y cells [American Type Culture Collection (ATCC), no. CRL-2266, RRID: CVCL_0019] were grown in RPMI 1640 (Gibco) medium completed with 10% fetal calf serum (FCS) and gentamicin (40 μg/ml), diluted 1:5 every 3rd/4th day. HEK293T cells were cultured in Dulbecco’s modified Eagle’s medium (DMEM; Eurobio) supplemented with 10% fetal bovine serum, 2 mM L-glutamine (Invitrogen), and penicillin and streptomycin (Invitrogen). All cells were cultured at 37°C under humidified conditions with 5% CO2.

### Zebrafish strain and husbandry

Zebrafish (*Danio rerio*) were raised and maintained as described [72]. Adult zebrafish were raised in 15 L tanks containing a maximum of 24 individuals, and under a 14 h to 10 h light-dark cycle. The water had a temperature of 28.5 °C and a conductivity of 200 µS and was continuously renewed. The fish were fed three times a day, with dry food and *Artemia salina* larvae. Embryos were raised in E3 medium, at 28.5 °C, under constant darkness. AB strain obtained from the Zebrafish International Resource Center (ZIRC) was used as wild-type for this study. For zebrafish embryos and larvae, both males and females were used since the sex can only be determined at 2 months of age. All animal experiments were carried out according to the guidelines of the Ethics Committee of IGBMC and ethical approval was obtained from the French Ministry of Higher Education and Research under the number APAFIS#49694-2024050316122698 .

### Mouse

Wild-type RjOrl:Swiss pregnant female mice (Janvier Labs, St Berthevin, France), used for generating E17.5 (Embryonic day 17,5) embryos for primary cultures of hippocampal neurons, and for harvesting P8 (postnatal day 8) pup brains, were housed at the animal house facility of the Institut de Génétique Moléculaire de Montpellier (France). Animals had a libitum access to food and water, with 12 h light-dark cycle. The mouse facility has been approved by the Département des pratiques de recherche réglementées “ Animaux à des fins scientifiques” – AFiS, under the approval number F3417216.

### Total cell extract preparations

To prepare the total cell extracts for native holdup assay, SH-SY5Y cells were seeded on 100 mM plates. When confluency was reached, the cells were washed with phosphate-buffered saline (PBS) once and collected by scraping with ice-cold lysis buffer [50 mM HEPES (pH 7.5), 150 mM NaCl, 1% Triton X-100, 1× cOmplete EDTA-free protease inhibitor cocktail, 5 mM TCEP, and 10% glycerol]. Lysates were sonicated 4 × 20 s with 1 s on /1 s/off cycle on ice. For mouse brain extracts, a P8 mouse pup was killed by decapitation and the brain was collected in ice-cold PBS and snap frozen as dry tissue. The brain was then homogenized in ice-cold lysis buffer [50 mM HEPES (pH 7.5), 150 mM NaCl, 1% Triton X-100, 1× cOmplete EDTA-free protease inhibitor cocktail, 5 mM TCEP, and 10% glycerol], using a Potter-Elvehjem homogenizer. For zebrafish lysates, 100 embryos (2 days post-fertilization; dpf) were anaesthetized with tricaine methanesulfonate and sodium bicarbonate. Then, the embryos were washed once in PBS and homogenized in 500 µl lysis buffer (1x PBS, 1% Triton X-100, 1× cOmplete EDTA-free protease inhibitor cocktail, 5 mM TCEP, and 10% glycerol). The homogenized lysates were sonicated 3x with 1-s-long pulses on ice. All lysates were incubated on roller shaker at 4 °C for 30 min. Then, lysates were centrifuged at 16 000 *xg* 4°C for 20 min, and the supernatant was kept for further analysis. Total protein concentration of the lysates from the SH-SY5Y cells as well as the mouse brain and the zebrafish embryos were measured by the standard Bradford method (Bio-Rad protein assay dye reagent, no. 5000006) using a bovine serum albumin (BSA) calibration curve (MP Biomedicals, no. 160069, diluted in lysis buffer) on a Bio-Rad SmartSpec 3000 spectrophotometer instrument. Lysates were diluted to 2 mg/ml total protein concentration and were snap-frozen in liquid nitrogen and stored at −80 °C until measurement.

To prepare cell extracts for coimmunoprecipitation and pull-down assay, HEK293T cells were lysed in lysis buffer [50 mM Tris-HCl pH 7.5, 150 mM NaCl, 1 mM EGTA, 1 mM EDTA, 0.5% (w/v) Triton X-100, 10% sucrose, 1 mM dithiothreitol (DTT), 1 mM Benzamidine, 0.5 mM AEBSF (4-(2-aminoethyl) benzenesulfonyl fluoride hydrochloride)] for 15 min, and lysates were cleared by centrifugation at 15, 000 xg for 10 min.

### Resin preparation and holdup experiment

To saturate streptavidin resin with biotinylated peptides (UBE3A 25-mer or biotin), 50 μl of streptavidin resin was mixed with biotin or peptide at 40 to 60 μM concentration in 6 to 6.5 resin volume (RV) for 60 min. To saturate streptavidin resin with biotinylated MBP, RLD2, UBE3A full-length proteins, 45 μl of streptavidin resin (Streptavidin Sepharose High Performance, Cytiva) was mixed with biotinylated MBP or MBP-RLD2 or MBP-UBE3A at 30 to 50 μM concentration in 20× resin volume for 60 min. After saturation, resins were washed once with 10 RV holdup buffer [50 mM Tris pH 7.5, 300 mM NaCl, and 1 mM TCEP, 0.22-μm filtered] and were depleted with biotin (1 RV, 5 min, holdup buffer supplemented with 1 mM biotin). Last, resins were washed four times with 10 RV holdup buffer.

Titration experiments were carried out by mixing control and bait-saturated resin and keeping the total resin volume constant as described [16]. Control and bait-saturated resins were prepared in larger amounts, and a serial dilution was prepared with these pre-saturated resins to achieve different resin ratios. The holdup mixture was incubated at 4 °C for 2 h. After incubation, the resin was separated from the supernatant by a brief centrifugation (15 s, 2 000 rpm). Then, half of the supernatant was removed by pipetting without delay to avoid any resin contamination. Holdup samples were mixed with 4× Laemmli buffer [120 mM Tris-HCl (pH 7.0), 8% SDS, 100 mM dithiothreitol, 32% glycerol, 0.004% bromophenol blue, and 1% β-ME] in a 3:1 ratio for further analysis.

### Co-immunoprecipitation assay

HEK293T cells were seeded on 100 mM plates and co-transfected using the JetPEI reagent (Polyplus) according to the manufacturer’s instructions, with pEGFP-RLD2 domain of HERC2 and pmCherry-DOCK10 (full-length), in their wildtype or mutant version, or control plasmids (pEGFP plasmid). 36 h post-transfection, cells were lysed as describe above. For each condition, GFP-RLD2 was immunoprecipitated from 1 mg of cleared total cell extracts by addition of 2 µg anti-GFP antibody (Torrey Pines Biolabs, TP401), coupled to 40 µl of Protein G-coated magnetic beads (Dynabeads Protein G, Invitrogen), and the co-precipitated mCherry-DOCK10 was detected by immunoblotting.

### RAC1 and CDC42 activation assays

The RAC1-GTP and CDC42-GTP pull-down assays were performed as described [14]. Briefly, HEK293T cells were transfected with pmCherry-DOCK10 plasmids, either wild-type or mutated, or with a control plasmid. 36 h post-transfection, the cells were lysed as described above, and processed for the pull-down assay (see below). For siRNA transfection, HEK293T cells were split and plated in 100 mm diameter dishes in the morning of Day 1. Late afternoon of Day 1, a pool of siCtrl (target sequences: UGGUUUACAUGUCGACUAA, UGGUUUACAUGUUGUGUGA, UGGUUUACAUGUUUUCUGA, UGGUUUACAUGUUUUCCUA from Dharmacon^TM^, Ref#: SO-2935020G) and a pool of siHERC2 (target sequences : GCACAGAGUAUCACAGGUA, CGAUGAAGGUUUGGUAUUU, GAUAAUACGACACAGCUAA, GCAGAUGUGUGCUAAGAUG from Dharmacon^TM^, Ref# : SO-3143778G) were transfected at a final concentration of 5 nM using 20 µl RNAiMAX lipofectamine^®^ (Life Technologies, Thermo Fisher Scientific) in Opti-MEM^®^ reduced serum medium (Thermo Fisher Scientific). In the morning of Day 2, the medium was changed, and pEGFP-DOCK10 and GFP control vector were transfected using JetPEI transfection reagent (Polyplus) according to the manufacturer’s instructions. On Day 4, cells were lysed and the RAC1-GTP and CDC42-GTP pull-downs were performed as described below.

The endogenous GTP-RAC1 (or GTP-CDC42) was pulled down by incubating the cleared lysates for 1 h at 4 °C with the CDC42/RAC1-interactive binding (CRIB) domain of PAK1 immobilized on GST-Sepharose beads (GE-Healthcare). Total HEK293T lysates and the corresponding pull-downs were processed for Western blotting. The level of GTP-bound (active) RAC1 (or CDC42) protein was assessed by band-intensity quantification using the Odyssey scanner from Li-COR Biosciences, and normalized to the total amount of RAC1 or CDC42 detected in the cell lysate.

### Immunoblotting

Native holdup samples were essentially processed as previously described [16]. Equal amounts of samples were loaded on 8 or 10% acrylamide gels. The transfer was done into PVDF membranes using a Trans-Blot Turbo Transfer System and a Trans-Blot Turbo RTA Transfer kit (Bio-Rad, no. 1704273). After 1 h of blocking in 5% milk, membranes were incubated overnight at 4 °C with primary antibody in 5% milk. The following antibodies and dilutions were used: anti-HERC2 (1:1000, BD Biosciences, no. 612366,RRID: AB_399728), anti-UBE3A (1:1000, Sigma-Aldrich, no. SAB1404508, clone 3E5, RRID: AB_10740376), anti–glyceraldehyde-3-phosphate dehydrogenase (GAPDH, 1:5000, Sigma-Aldrich azide-free version of AB_2924240), anti-alpha tubulin (1:5000, Sigma-Aldrich, no.T9026, clone DM1A, RRID:AB_477593), anti-DOCK10 (1:500, Sigma-Aldrich, no. HPA058106, RRID: AB_2683606), anti-ARID4A (1:1000, CST, no. 97780) and anti-PCM1 (1:250, CST, no. 5213, RRID: AB_10556960). Membranes were washed three times with Tris-buffered saline (TBS)–Tween and incubated with secondary antibody [Jackson ImmunoResearch, peroxidase-conjugated Affinipure goat anti-mouse (H + L), no. 115-035-146 RRID: AB_2307392 and goat anti-rabbit (H + L), no. 111-035-003 RRID: AB_2313567] at room temperature for 1 h in 5% milk (dilution 1:10,000). After washing three times with TBS-Tween, membranes were exposed to chemiluminescent horseradish peroxidase substrate (Immobilon, no. WBKLS0100 or SuperSignal™ West Atto Ultimate Sensitivity Substrate, no. 38554) and visualized in a docking system (Amersham ImageQuant™ 800, GE). Densitometry analysis was carried out on raw Tif images by using Fiji ImageJ 1.53c. Between different primary antibody labeling, the membranes were either exposed to 15% (v/v) H_2_O_2_ to remove secondary signal (in the case of different species) or stripped with mild stripping buffer [glycine (15 g/liter), SDS (1 g/liter), and 1% Tween 20 (pH 2.2)] to remove primary signal (in the case of same species). All holdup titrations were run and western-blotted in triplicate, leading to three distinct membranes that were subjected to densitometry analysis. This allowed us to perform statistical analysis, warranting the highest precision and accuracy for the determination of binding constants. For each holdup titration presented in the main text of the article, the three corresponding blotted membranes are provided in supplemental figures.

For coimmunoprecipitation and pull down, the following antibodies and dilution were used: anti-mCherry (1:1000, Invitrogen, no. PA5-34974, RRID: AB_2552323), anti-GFP- (1:3000, Torrey Pines Biolabs, no. TP401, RRID: AB_2313770), anti-RAC1 (1:1000, BD Biosciences, n o . 610650, RRID: AB_397977), anti-CDC42 (1:250, BD Biosciences, no. 610929, RRID: AB_398244). Immunoblot detections and band-intensity quantifications were made with the Odyssey scanner from Li-COR Biosciences.

### Primary culture and transfection of hippocampal neurons

Neuronal hippocampal cultures were prepared from embryonic day 17.5 RjOrl: Swiss embryos, as described (Bonnet et al, 2023). Neurons were transfected at 8 DIV (Days *In Vitro*) with cDNA plasmid constructs of pmCherry-DOCK10, wildtype or mutated, or a control plasmid, using the Lipofectamine 2000 reagent (ThermoFisher), according to the manufacturer’s instructions. At DIV11, neurons were fixed with 4% paraformaldehyde/4% sucrose in PBS for 10 min and processed for microscopic examination.

### Immunolabelling and dendritic spine quantification

Fixed neurons were permeabilized in 0.15% Triton-X-100/PBS for 3 min. Immunostaining was performed with GFP antibody (Aves Labs, no. GFP-1020, RRID: AB_10000240) and mCherry (Invitrogen, PA5-34974, RRID: AB_2552323) and the corresponding Alexa Fluor 488 and 546-conjugated secondary antibodies (ThermoFisher Scientific). All coverslips were mounted in Mowiol reagent (Sigma Aldrich). Images were acquired using a fluorescence microscope (Zeiss Axioimager Z2) and a 63x objective (NA 0.8). Morphometric analyses were performed in different fields from at least three different cultures, using the ImageJ software (National Institutes of Health, Bethesda, MD). Spines were defined as dendritic protrusions with a neck and a head. Dendrites were randomly selected and spines were manually counted over a 50-µM length of dendrite. Data were then expressed as density of spines/10- µm length of dendrite. All experiments were conducted in a blinded manner, and dendritic spines were analyzed from at least 20 neurons from at least three independent cultures.

## Acknowledgments

We thank André Mitschler, Arantxa Agote-Arán and Izabela Sumara for insightful scientific discussions and for providing preliminary results. We thank the cell culture platform of the IGBMC for their help in cell culturing. We thank Sandrine Geschier from the IGBMC zebrafish facility. We thank Pierre Poussin-Courmontagne for the support in protein crystallization and the staff at SOLEIL and SLS for the help and assistance during data collection. The project was supported by the Ligue contre le cancer (équipe labellisée 2015 to GT), the Agence Nationale de la Recherche (grant ANR-18-CE92-0017, ANR-22-CE44-0018, and ANR-22-CE11-0026 to GT, ANR-19-CE16-0004 to Anne Debant), Dup15q Alliance/Angelman Syndrome Foundation joined Award (2019-2021 to GT). As a member of the IGBMC institute, we benefited from the French Infrastructure for Integrated Structural Biology (FRISBI) ANR-10-INSB-05–01, from Instruct-ERIC, from IdEx Unistra (ANR-10-IDEX-0002), from SFRI-STRAT’US project (ANR 20-SFRI-0012), and from EUR IMCBio (ANR-17-EURE-0023) under the framework of the French Investments for the Future Program as a member of the Interdisciplinary Thematic Institute IMCBio, as part of the ITI 2021–2028 program of the University of Strasbourg, CNRS and Inserm. We also acknowledge the imaging facility MRI (Montpellier, France), member of the national infrastructure France-BioImaging supported by the French National Research Agency (ANR-10-INBS-04, «Investments for the future»). We are grateful to the Réseaux des Animaleries de Montpellier (RAM-ZEFI) for animal care and maintenance. The Fondation pour la recherche médicale (FRM) supported AD (prix de thèse Pomaret-Delalande sur les maladies rares, 2019-2021) and JWL (SPF202309017619), and Anne Debant (équipe FRM EQU202403018054, 2024-2027).

