## Supplementary information for "Structural analysis of HERC2/UBE3A and HERC2/DOCK10 complexes provides new insights into the molecular basis of Angelman, Angelman-like and Dup15q Syndromes"

&: equal contributors

*: corresponding authors.

**Supplementary Figure 1.** The results of the three native holdup titrations with UBE3A full-length protein using SH-SY5Y cell lysates used for the apparent affinity determination. The protein levels of full-length endogenous HERC2 and α-tubulin were visualized by Western-blot. All membranes were stripped with stripping buffer (15 g/L glycine, 1 g/L SDS, 1% v/v Tween20, pH 2.2) before being re-probed with different antibodies as indicated. Membrane 1 was used in Figure 1A.

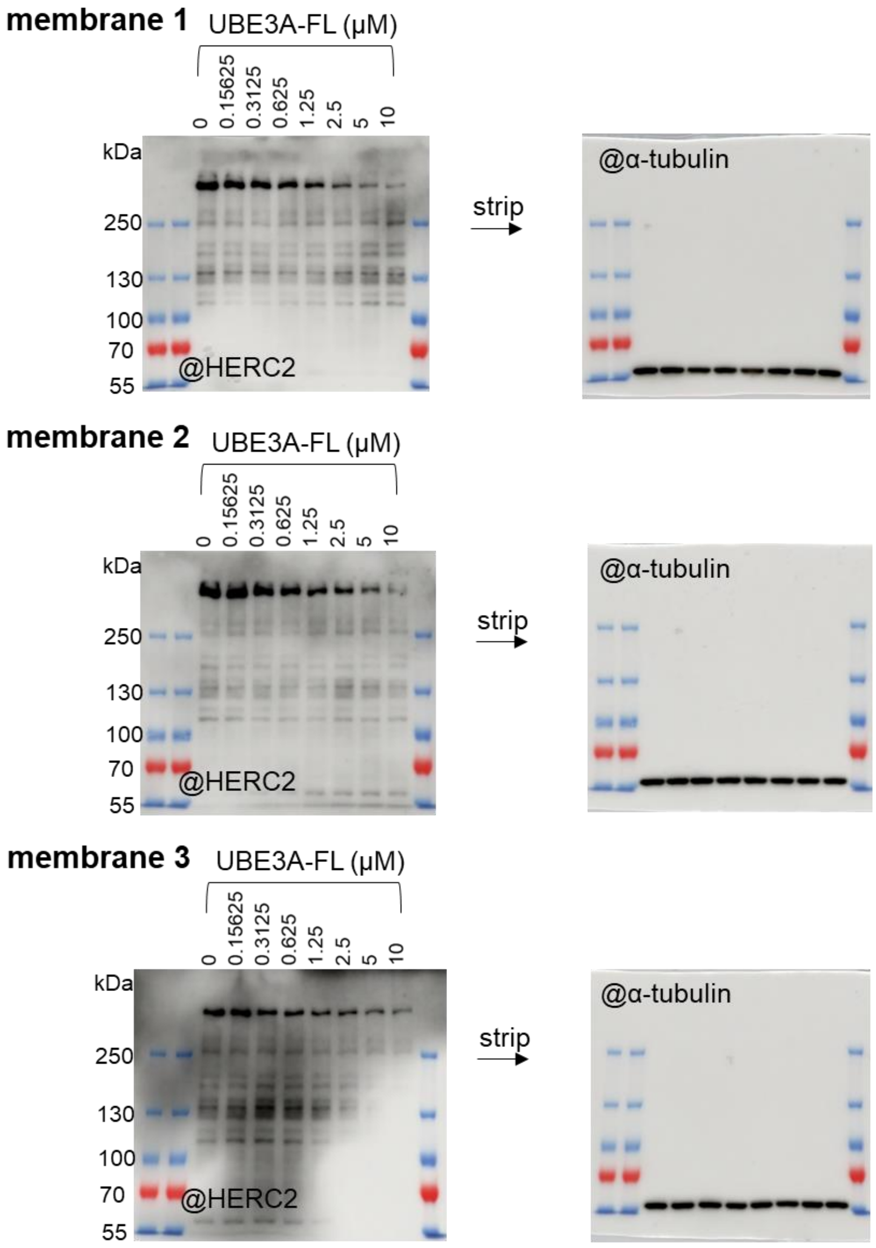

**Supplementary Figure 2.** The results of the three native holdup titrations with UBE3A 150-174 using SH-SY5Y cell lysates used for the apparent affinity determination. The protein levels of full-length endogenous HERC2 and α-tubulin were visualized by Western-blot. All membranes were stripped with stripping buffer (15 g/L glycine, 1 g/L SDS, 1% v/v Tween20, pH 2.2) before being re-probed with different antibodies as indicated. Membrane 1 and 2 were used in Figure 1C.

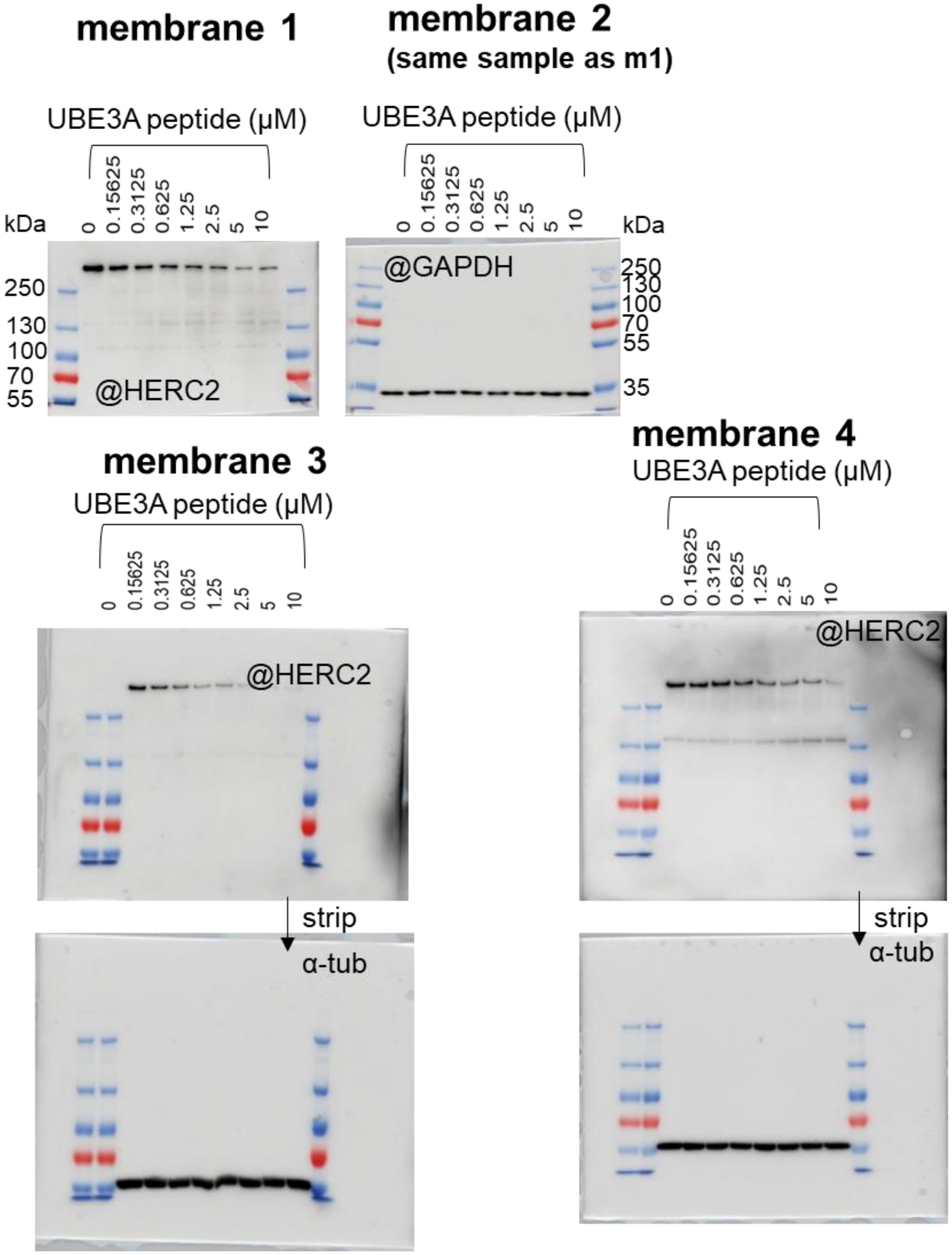

**Supplementary Figure 3.** Binding affinities of the RLD2 to the recombinant UBE3A full-length proteins or the different peptides derived from UBE3A residues 150 – 200 obtained by fluorescence polarization (FP). A. The direct binding curves of purified HERC2 RLD2 domain with fluorescein labeled UBE3A 180-197 (fluo-UBE3Apep) were monitored in fluorescence polarization by titrating 50 nM fluo-UBE3Apep with an increasing amount of HERC2 RLD2 domain (0 – 15 μM). The clear increase in FP indicates a binding event has occurred**.** B-L. The reversibility of the complex formation was monitored with a competitive measurement by titrating the RLD2/fluo-UBE3Apep complex with an increasing amount of non-labeled UBE3A full-length proteins or different UBE3A peptides as indicated. A decrease in the FP signal indicates the reversible complex formation. Concluding from the competition measurement, peptides of UBE3A 150-174 (C) showed strongest affinity to RLD2 followed by UBE3A full-length proteins (B), UBE3A 150-200 (D) and UBE3A 160-174 (G) that formed complex at a similar affinity range. As compared to full-length UBE3A proteins, the peptides of UBE3A 150-164 (E), UBE3A 155-169 (F), UBE3A 165-179 (H) formed complex at 20 - to 200-fold lower affinity whereas UBE3A 170-184 (I), UBE3A 175-189 (J), UBE3A 180-194 (K) and UBE3A 185-200 (L) show no significant decrease in FP signals indicating irreversible complex formation thus no binding to RLD2 domain.

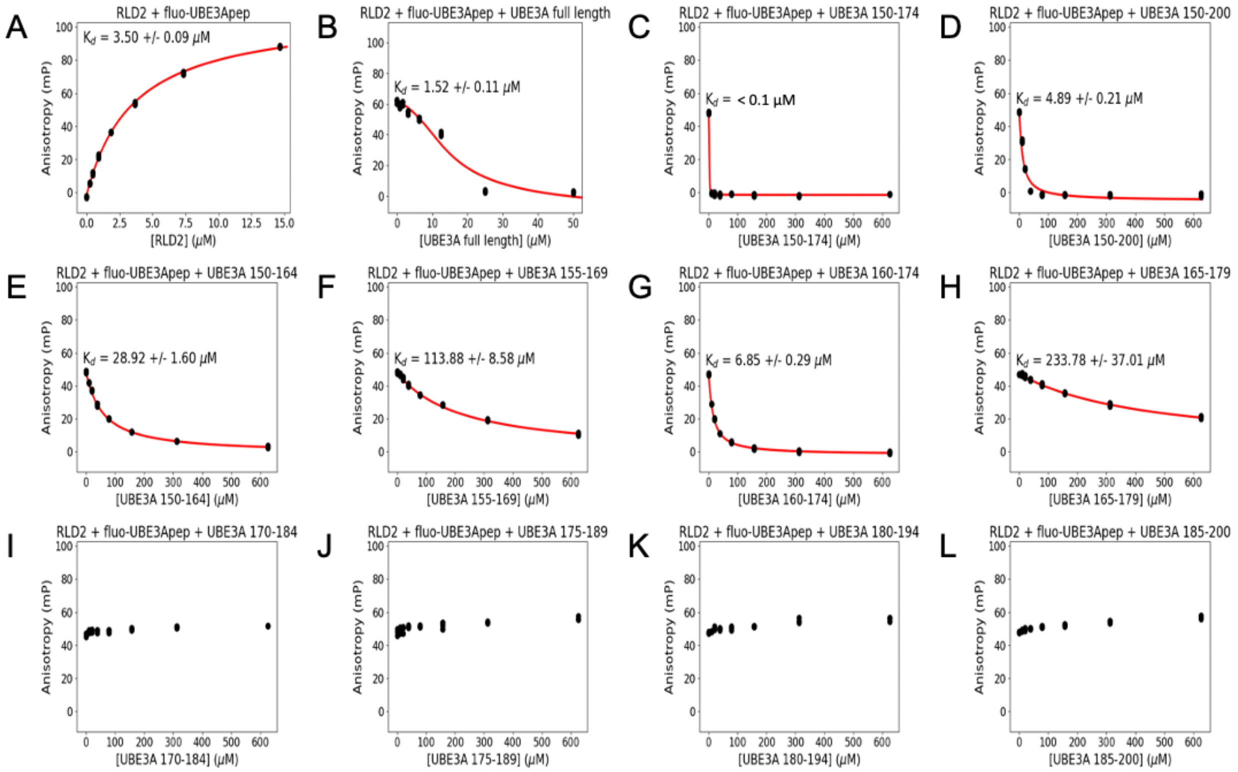

**Supplementary Table 1.** The summary of the binding affinities of the RLD2 to the recombinant UBE3A full-length protein or the different peptides derived from UBE3A residues 150 – 200 obtained by competitive fluorescence polarization (FP). The numbering of the UBE3A residues is based on the numbering of UBE3A isoform I.

| Proteins | Peptide Name | Sequences | KD (μM) |
|---|---|---|---|
| UBE3A | UBE3A full-length | UniProt Q05086-2 | 1.52 ± 0.11 |
| | UBE3A 150-174 | HTKEELKSLQAKDEDKDEDEKEKAA | < 0.1 |
| | UBE3A 150-200 | HTKEELKSLQAKDEDKDEDEKEKAAC SAAAMEEDSEASSSRIGDSSQGDNN | 4.89 ± 0.21 |
| | UBE3A 150-164 | HTKEELKSLQAKDED | 28.92 ± 1.60 |
| | UBE3A 155-169 | LKSLQAKDEDKDEDE | 113.88 ± 8.58 |
| | UBE3A 160-174 | AKDEDKDEDEKEKAA | 6.85 ± 0.29 |
| | UBE3A 165-179 | KDEDEKEKAACSAAA | 223.78 ± 37.01 |
| | UBE3A 170-184 | KEKAACSAAAMEEDS | No binding |
| | UBE3A 175-189 | CSAAAMEEDSEASSS | No binding |
| | UBE3A 180-194 | MEEDSEASSSRIGDS | No binding |
| | UBE3A 185-200 | EASSSRIGDSSQGDNN | No binding |

**Supplementary Table 2** Data collection and refinement parameters.

| PDB | 7Q40 | 7Q41 | 7Q42 | 7Q43 | 7Q44 | 7Q45 | 7Q46 |
|---|---|---|---|---|---|---|---|
| Resolution (last shell) (Å) | 54.27-2.35 (2.41-2.35) | 48.72-3.01 (3.20-3.01) | 49.29-1.95 (2.00-1.95) | 49.67-2.40 (2.46-2.40) | 49.52-2.20 (2.26-2.20) | 48.47-2.10 (2.15-2.10) | 49.16-2.46 (2.52-2.46) |
| Space group | $P4_32_12$ | $P4_32_12$ | $P4_32_12$ | $P4_32_12$ | $P4_32_12$ | $P4_32_12$ | $P4_32_12$ |
| Cell parameters | a=b=108.54Å c=243.09 Å α=β=γ=90° | a=b=108.952Å c=243.16Å α=β=γ=90° | a=b=108.952Å c=241.56Å α=β=γ=90° | a=b=108.82Å c=243.21 Å α=β=γ=90° | a=b=108.46Å c=242.85 Å α=β=γ=90° | a=b=108.39Å c=242.31 Å α=β=γ=90° | a=b=107.66Å c=241.21 Å α=β=γ=90° |
| Completeness (last shell) (%) | 98.8 (91.0) | 99.2 (97.7) | 99.9 (99.3) | 99.7 (100) | 99.9 (99.7) | 99.2 (98.6) | 100 (99.9) |
| Redundancy (last shell) | 26.7(21.6) | 5.9 (6.1) | 27.1 (24.3) | 37.7 (34.7) | 26.7 (26.7) | 26.8 (27.0) | 26.5 (27.9) |
| I/σ(I) (last shell) | 13.21 (0.93) | 5.81 (1.05) | 10.93 (1.03) | 5.77 (1.00) | 8.65 (1.25) | 9.67 (1.16) | 6.33 (1.20) |
| CC(1/2) (last shell) (%) | 99.8 (47.3) | 96.8 (40.1) | 99.8 (61.0) | 98.3 (41.0) | 99.4 (44.4) | 99.5 (45.6) | 98.4 (49.8) |
| Rpim (last shell) (%) | 4.8 (52.3) | 16.3 (85.0) | 5.2 (43.3) | 18.4 (78.8) | 9.0 (63.3) | 7.9 (65.2) | 14.1 (72.6) |
| $R_{work}/R_{free}$ | 17.9/23.0 | 19.8/25.5 | 16.8/20.6 | 20.2/26.5 | 18.2/23.1 | 18.4/22.4 | 17.9/24.5 |
| RMSD Bond (Å) | 0.007 | 0.009 | 0.011 | 0.008 | 0.003 | 0.004 | 0.007 |
| RMSD Angle (°) | 0.854 | 1.094 | 1.134 | 0.911 | 0.634 | 0.788 | 0.892 |
| Ramachandran favored (%) | 97 | 96 | 97 | 97 | 98 | 97 | 97 |
| Ramachandran allowed (%) | 3 | 4 | 3 | 3 | 2 | 3 | 3 |
| Ramachandran outliers (%) | 0 | 0 | 0 | 0 | 0 | 0 | 0 |
| Average B-factor ($Å^2$) | 56.0 | 58.0 | 33.0 | 38.0 | 37.0 | 35.0 | 36.0 |

**Supplementary Table 3.** DxDKDxD motif-containing proteins identified by SlimSearch.

| Proteins | Sequences | Start | End | Remarks |
|---|---|---|---|---|
| PCM1 | ageidDEDKDKDetetv | 1737 | 1743 | Proteins with perfect and accessible DxDKDxD motif |
| ARID4A | ikkqeDSDKDSDeeeeK | 537 | 543 | |
| USP35 | wgrgfDEDKDEDegspg | 990 | 996 | |
| MYT1 | vevrsDDDKDEDthsrk | 349 | 355 | |
| DOCK10 | eidheDADKDEDttshs | 158 | 164 | |
| ARIP4 | fssnDDEDKDDDvievt | 1454 | 1460 | |
| UBE3A | slqakDEDKDEDekeka | 185 | 191 | |
| BAZ2B | seeedDDDKDQDesdsd | 651 | 657 | |
| SMC6 | qeeleDFDKDGDedeck | 25 | 31 | |
| MED1 | kdrdrDRDKDRDkkksh | 1515 | 1521 | |
| RERE | mtaDKDKDKDkekdr | 4 | 10 | |
| RCN2 | drfvnDYDKDNDgrldp | 238 | 244 | Proteins with perfect but inaccessible DxDKDxD motif |
| CALR | edkedDEDKDEDeedee | 388 | 394 | |
| MIGA1 | nsladDIDKDTDitmkg | 297 | 303 | |

**Supplementary Figure 4.** The results of the three native holdup titrations with MBP-RLD2 using mouse brain lysates used for the apparent affinity determination. The protein levels of full-length endogenous UBE3A and α-tubulin were visualized by Western-blot. All membranes were cut between 72 -130 kDa marker band to incubate with anti-UBE3A while membrane cut between 35-72 kDa marker were incubated with anti-α-tubulin. Membrane 3 was used in Figure 5C.

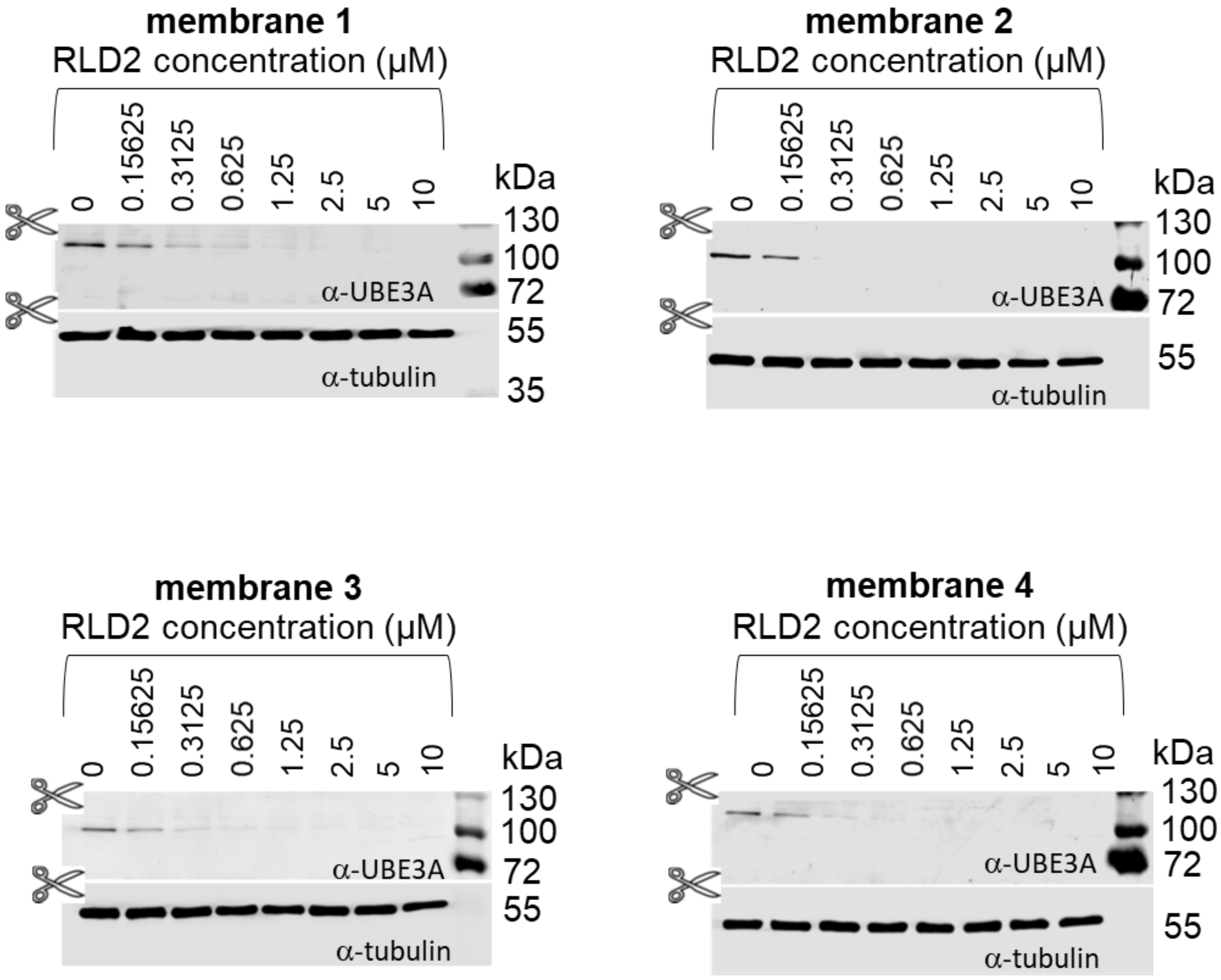

**Supplementary Figure 5.** The results of the three native holdup titrations with MBP-RLD2 using zebrafish embryos lysates used for the apparent affinity determination. The protein levels of full-length endogenous UBE3A and α-tubulin were visualized by Western-blot. All membranes were stripped with stripping buffer (15 g/L glycine, 1 g/L SDS, 1% v/v Tween20, pH 2.2) before being re-probed with different antibodies as indicated. Membrane 3 was used in Figure 5D.

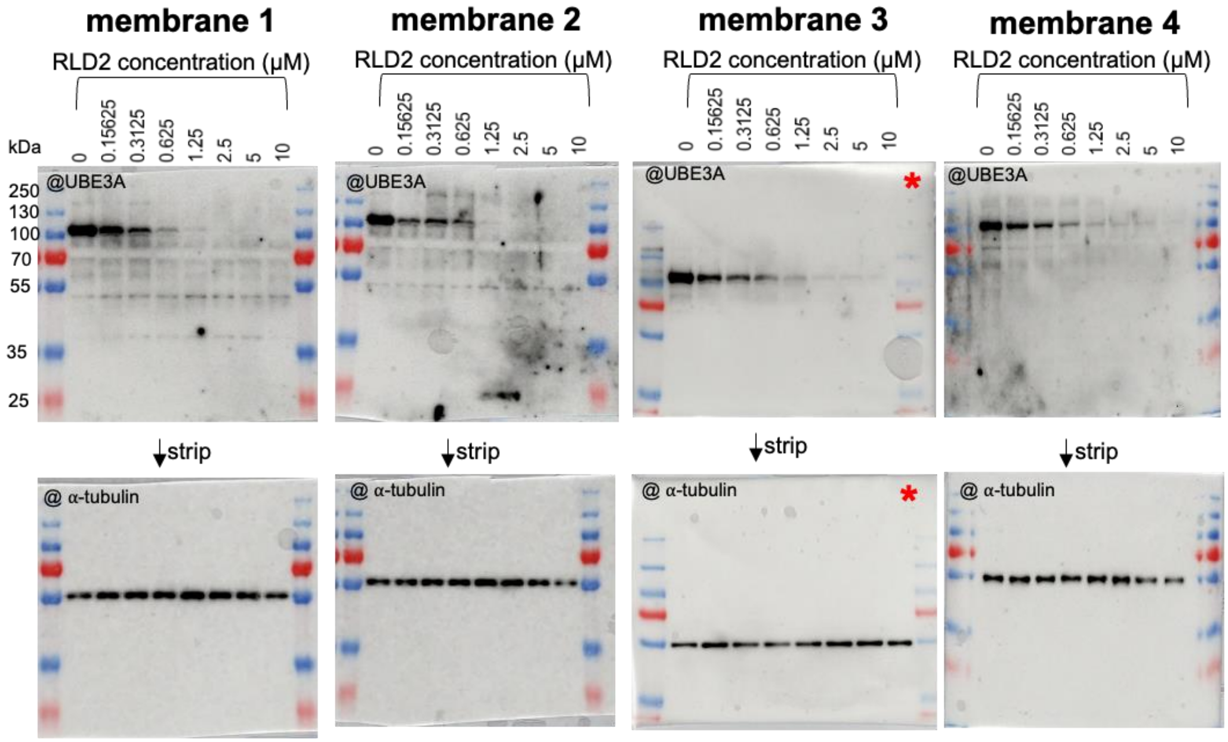

**Supplementary Figure 6.** Binding affinities of the RLD2 to the different peptides derived from the different DxDKDxD-containing proteins (listed in Supplementary Table 4) by competitive fluorescence polarization (FP). The reversibility of the complex formation was monitored with a competitive measurement by titrating the RLD2/fluo-UBE3Apep complex with an increasing amount of non-labeled peptides as indicated. A decrease in the FP signal indicates the reversible complex formation. According to Alphafold prediction, the DxDKDxD motif of RCN2, CALR and MIGA1 are inaccessible. Hence, the peptides are not measured for competitive FP.

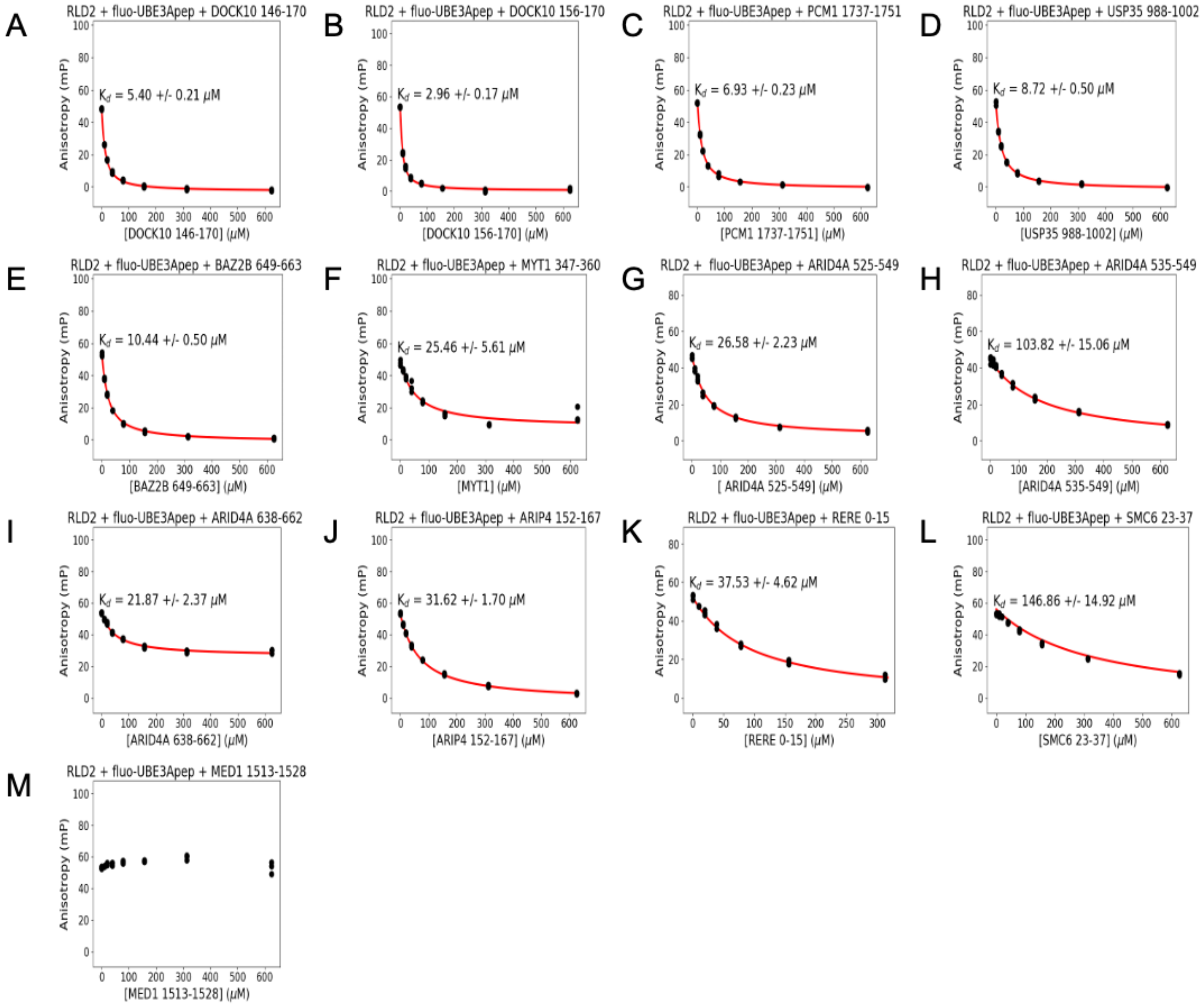

**Supplementary Table 4.** The summary of the binding affinities of the RLD2 to the different peptides derived from the different DxDKDxD-containing proteins by competitive fluorescence polarization (FP). The numbering of the residues of each protein is based on the numbering of the canonical sequence.

| Proteins | Peptide Name | Sequences | KD (μM) |
|---|---|---|---|
| DOCK10 | DOCK10 146-170 | KLPSHSFEIDHEDADKDEDTTSHSS | 5.40 ± 0.21 |
| | DOCK10 156-170 | HEDADKDEDTTSHSS | 2.96 ± 0.17 |
| PCM1 | PCM1 1737-1751 | DEDKDKDETETVKQT | 6.93 ± 0.23 |
| USP35 | USP35 988-1002 | GFDEDKDEDEGSPGG | 8.72 ± 0.50 |
| BAZ2B | BAZ2B 649-663 | EDDDDKDQDESDSDT | 10.44 ± 0.50 |
| MYT1 | MYT1 347-360 | RSDDDKDEDTHSRK | 26.58 ± 2.23 |
| ARID4A | ARID4A 525-549 | PKQKEKKIKKQEDSDKDSDEEEEKS | 103.82 ± 15.06 |
| | ARID4A 535-549 | QEDSDKDSDEEEEKS | 103.82 ± 15.06 |
| | ARID4A 638-662 | KKQKKKAKNKEDSEKDEKRDEERQ | 21.87 ± 2.37 |
| ARIP4 | ARIP4 1452-1467 | NDDEDKDDDVIEVTGK | 31.62 ± 1.70 |
| RERE | RERE 0-15 | MTADKDKDKDKEKDR | 37.53 ± 4.62 |
| SMC6 | SMC6 23-37 | LEDFDKDGDEDECKG | 146.86 ± 14.92 |
| MED1 | MED1 1513-1528 | DRDRDKDRDKKKSHS | No binding |

**Supplementary Figure 7.** The results of the three native holdup titrations with MBP-RLD2 using SH-SY5Y lysates used for the apparent affinity determination. The protein levels of full-length endogenous UBE3A, DOCK10, PCM1, ARID4A and GAPDH were visualized by Western-blot. All membranes were stripped with 15% v/v hydrogen peroxide ($H_2O_2$) or stripping buffer (15 g/L glycine, 1 g/L SDS, 1% v/v Tween20, pH 2.2) before re-probed with different antibodies as indicated. Membranes used in Figure 2B and 6C are indicated with red asterisk (*).

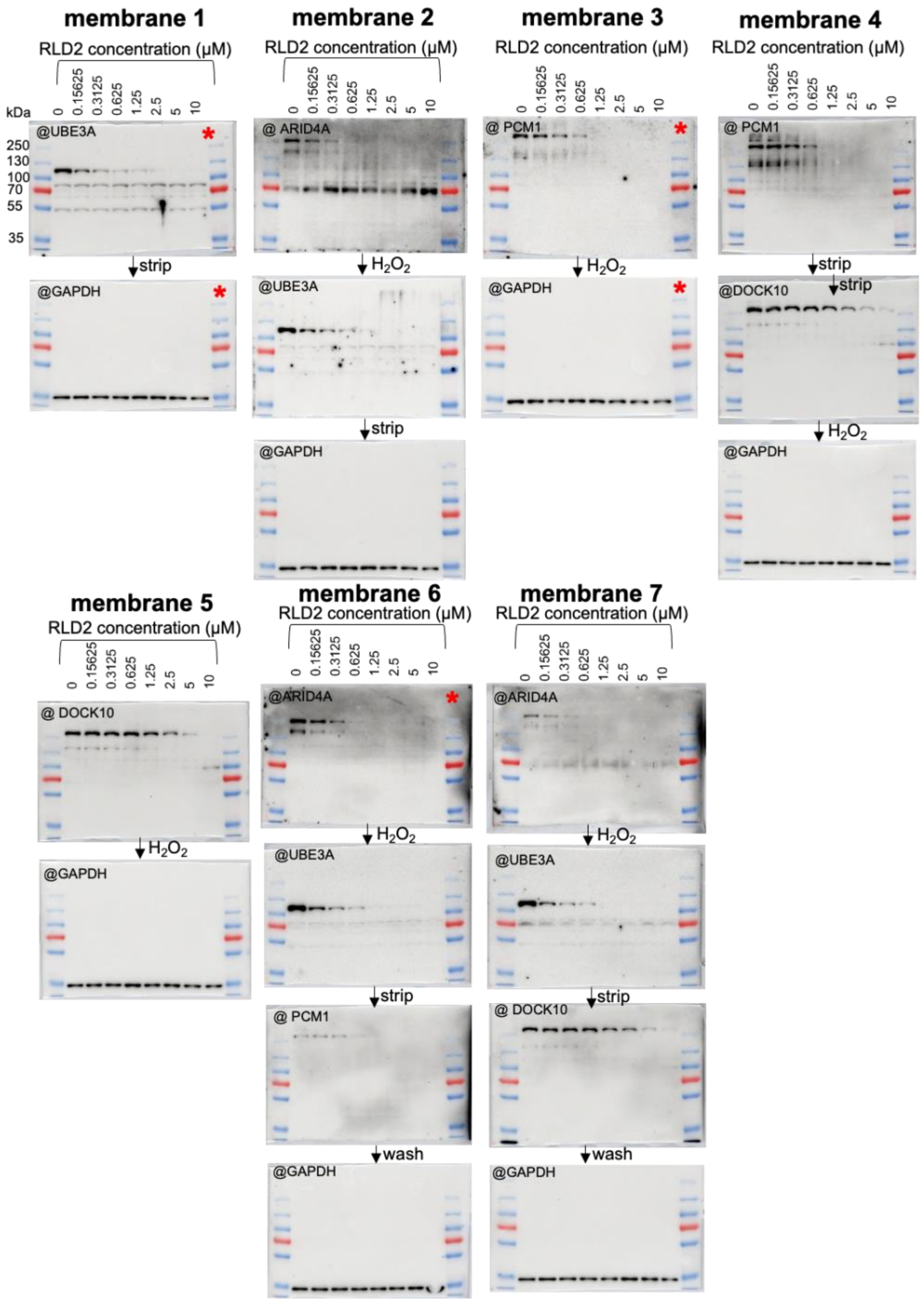

membrane 1
RLD2 concentration (μM)
0
0.15625
0.3125
0.625
1.25
2.5
5
10
kDa
250
130
100
70
55
35
@UBE3A
strip
@GAPDH
membrane 2
RLD2 concentration (μM)
@ ARID4A
$H_2O_2$
@UBE3A
strip
@GAPDH
membrane 3
RLD2 concentration (μM)
@ PCM1
$H_2O_2$
@GAPDH
membrane 4
RLD2 concentration (μM)
@ PCM1
strip
@DOCK10
strip
$H_2O_2$
@GAPDH
membrane 5
RLD2 concentration (μM)
@ DOCK10
$H_2O_2$
@GAPDH
membrane 6
RLD2 concentration (μM)
@ARID4A
$H_2O_2$
@UBE3A
strip
@ PCM1
wash
@GAPDH
membrane 7
RLD2 concentration (μM)
@ARID4A
$H_2O_2$
@UBE3A
strip
@ DOCK10
wash
@GAPDH

**Supplementary Figure 8.** The results of the native holdup titrations with MBP-RLD2 using SH-SY5Y lysates used for the apparent affinity determination. The protein levels of full-length endogenous MYT1, α-tubulin and GAPDH were visualized by Western-blot. It should be noted that due to the highly intrinsically disordered region and acidic properties of MYT1, it migrates anomalously at higher molecular weight on SDS-PAGE. All membranes were stripped with 15% v/v hydrogen peroxide ($H_2O_2$) before re-probed with different antibodies as indicated. Membranes used in Figure 6C are indicated with red asterisk (*).

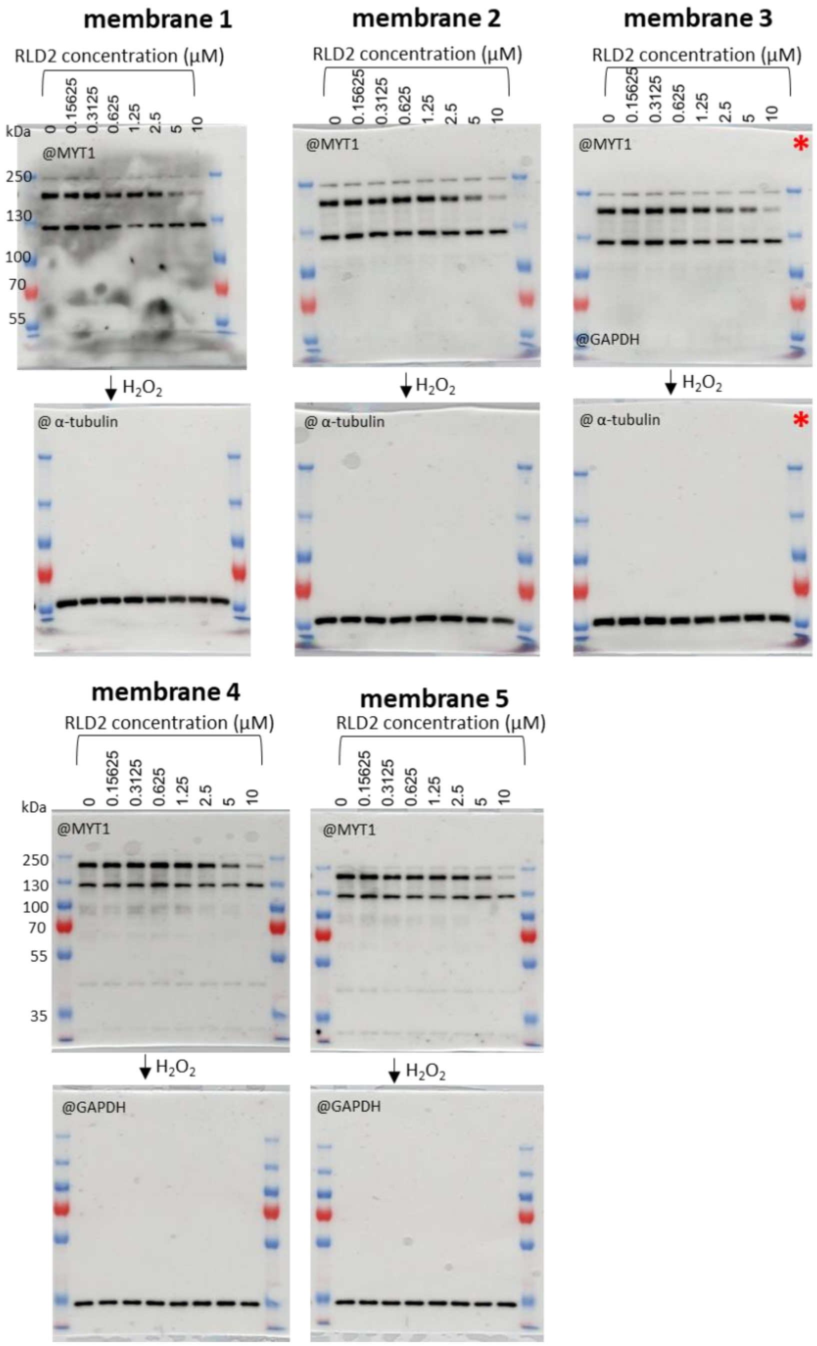

**Supplementary Figure 9**. Full immunoblot of the immunoprecipitation assay (IP) shown in Figure 7A. The immunoblot of the Input (total lysates) is shown in full in Figure 7A.

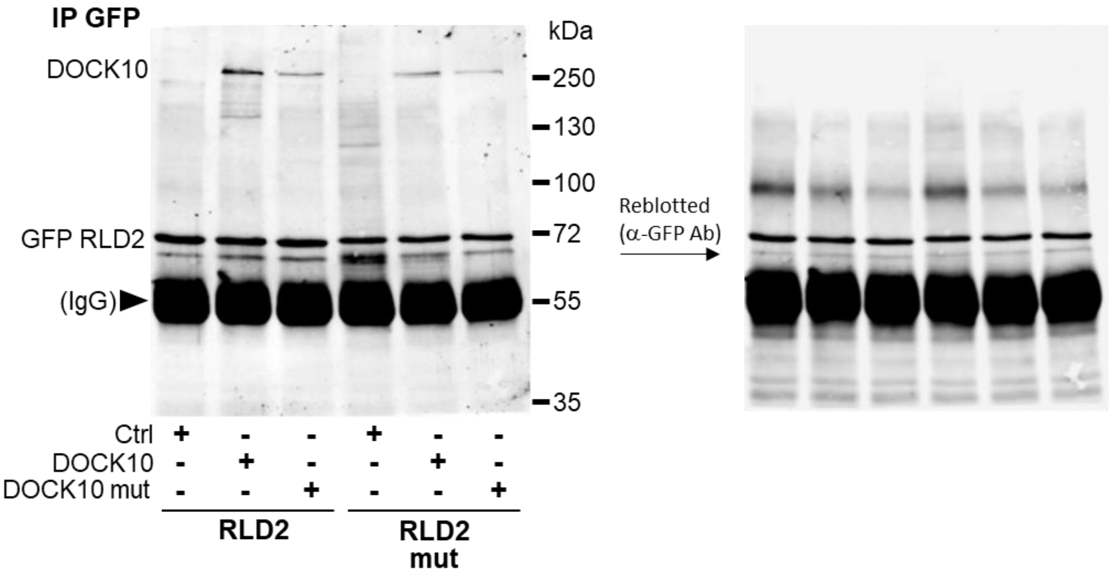

**Supplementary Figure 10**. **A-D.** Full immunoblots of the GTPase-pulldown assays presented in Figure 7B-D. Panel A corresponds to full blots of Figure 7B, CDC42 pulldown and DOCK10 expression blot. Panel B corresponds to full blots of Figure 7C, RAC1 pulldown and DOCK10 expression blot. Panels C correspond to full blots of the CDC42 and RAC1 pulldown of Figure 7D, and panel D are the correlated full expression blots, detected with DOCK10 antibody (left panel) and HERC2 antibody (right panel). **E.** TRIO-mediated RAC1-GTP pull down assays performed in the absence of HERC2. HEK293T cells were co-transfected with siHERC2 or siCtrl and pEGFP-TRIO or a control vector (siRNA transfected for 72hrs, cDNA transfected for 48hrs). Active GTP-RAC1 were affinity purified as described. Purified GTP-bound and total RAC1 were detected by Western blot, using the relevant anti-RAC1 antibodies. Protein expression of TRIO and HERC2 in the cell lysates was verified by immunoblotting with an anti-GFP and anti-HERC2 antibody. The % of RAC1 activation (right hand histogram) was calculated from at least 4 independent experiments (mean ±SEM). Statistical analyses were made using the non-parametric two-tailed Mann-Whitney test (*p<0.05). **F**. Full immunoblots of the GTPase-pulldown assays and expression immunoblots presented in E.

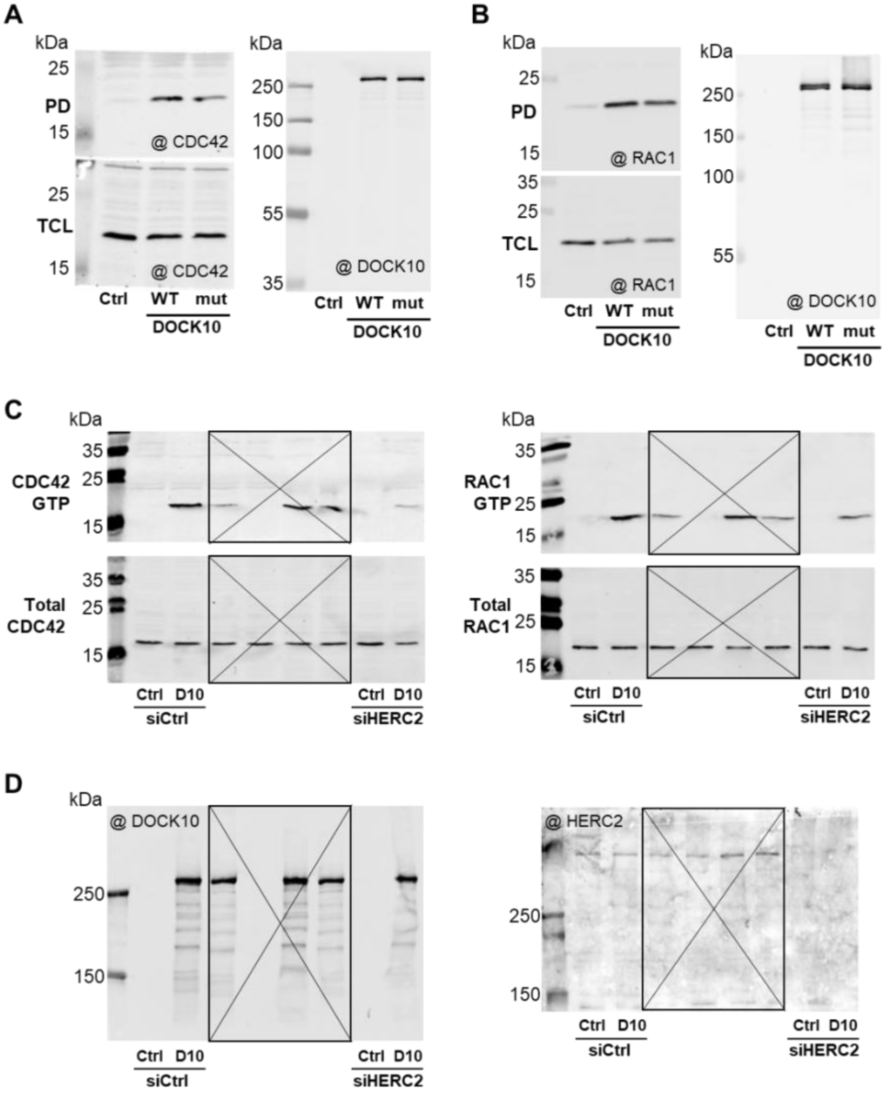

**E**

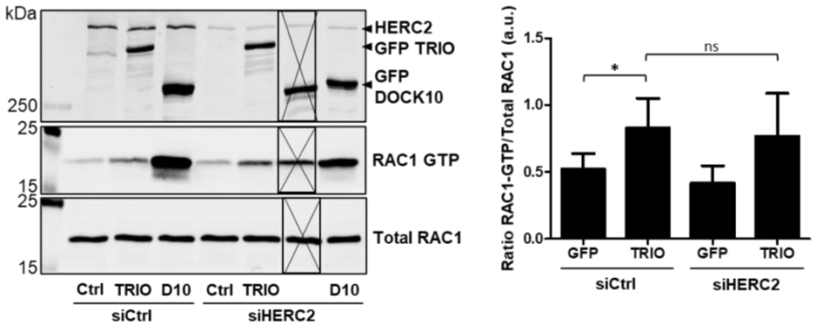
kDa
250
25
15
25
15
HERC2
GFP TRIO
GFP DOCK10
RAC1 GTP
Total RAC1
Ctrl TRIO D10 Ctrl TRIO D10
siCtrl
siHERC2
Ratio RAC1-GTP/Total RAC1 (a.u.)
1.5
1.0
0.5
0.0
*
ns
GFP TRIO GFP TRIO
siCtrl
siHERC2

**F**

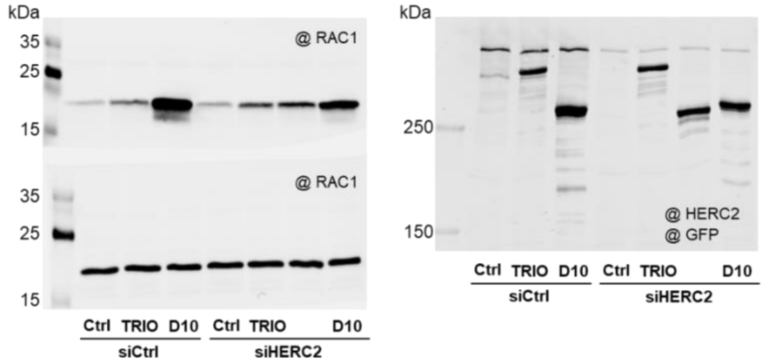
kDa
35
25
15
@ RAC1
35
25
15
@ RAC1
Ctrl TRIO D10 Ctrl TRIO D10
siCtrl
siHERC2
kDa
250
150
@ HERC2
@ GFP
Ctrl TRIO D10 Ctrl TRIO D10
siCtrl
siHERC2

**Supplementary Figure 11. Predicted structure of Apo-DOCK10 and experimental structures of DOCK2/ELMO1 and DOCK5/ELMO1 protomers.** (**A**) The main folded region of predicted DOCK10 comprises a DHR1-C2/armadillo solenoid/DHR2 arrangement reminiscent of that observed in DOCK2 and 5. Furthermore, the PH domain of DOCK10 is spatially proximal to the DHR2 domain of DOCK10, comparably to the proximity of the PH domain of ELMO1 to the DHR2 domains of DOCK2 and 5. (**B-C**) The DOCK2/ELMO1 and DOCK5/ELMO1 protomers correspond to experimentally determined Cryo-EM structures [1, 2]. DOCK2 and DOCK5 are colored blue to red from N-term to C-term. The ELMO obligate partner of DOCK2 and 5 is colored pale blue, except its PH domain which is colored blue.

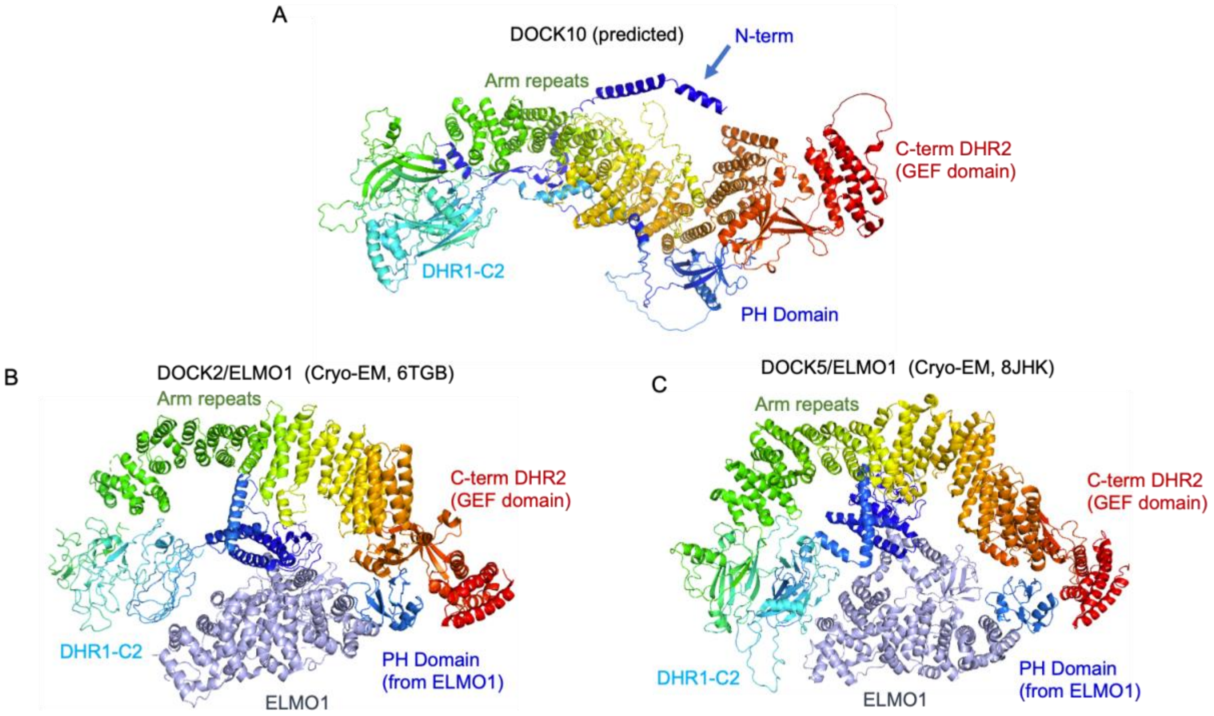

**Supplementary Figure 12. The N-terminal extension of DOCK10, shown in three distinct orientations.** The long N-terminal extension (dark blue to lighter blue, cartoon representation) is consistently predicted as a precisely positioned structure that wraps around the surface of the main folded unit composed of the DHR1-C2 armadillo body and DHR2 domain (cyan to red, spheres representation). It includes a kinked $\alpha$-helix at the extreme N-terminus, an $\alpha\beta\alpha\beta\alpha$ structure that extends towards the DHR1-C2 domain, the RLD2-binding DxDKDxD motif, and a PH domain contiguous to the DHR2 domain.

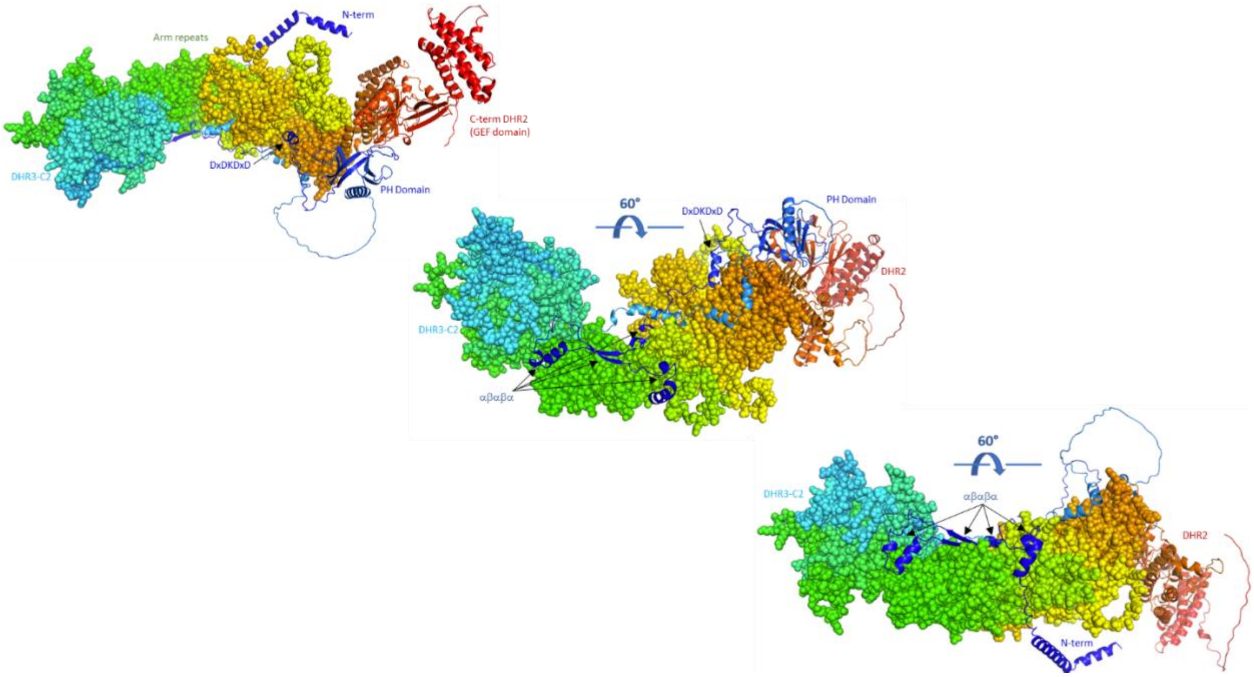

**Supplementary Figure 13: A. Alignment of extreme N-terminal sequences of human DOCK10, 11 and 9.** In DOCK10, the N-terminal sequence RTRRFTRSL forms a capping motif that interacts with Rac1 and CDC42 RhoGTPases in predicted DOCK10/Rac1 and DOCK10/CDC42 complexes. The residues directly engaged in the interface with the GTPase in these complexes are colored red in the alignment. They are strictly conserved in DOCK10, 9 and 11, indicating that the N-terminal helix of all DOCKD proteins is able to cap the bound RhoGTPase in a similar manner. **B. Zoomed view of the predicted full-length DOCK10/CDC42 complex showing details of the GDP-GTP exchange mechanism.** The CDC42 protein (pale blue) is represented in the same orientation as in the Yang et al., [3]. The GTP molecule (green) and the Mg2+ ion (blue) come from superimposition with the X-ray structure of GTP-bound DHR2 domain of DOCK9 (2wmo.pdb). As observed in the Yang et al article [3], Mg2+ expulsion is carried out by a conserved Val residue provided by the DHR2 domain. The N-terminal cap of DOCK10 (dark blue) establishes contacts with switch 2 of CDC42 (purple). This probably helps to rigidify switch 2 and keep Ala 59 of CDC42 (purple spheres) away from the Mg2+ ion. Ala 59 can serve as a Mg2+ expulser when CDC42 is bound to other GEFs [3], while Mg2+ expulsion by DOCK GEFs is performed by insertion of a conserved Val residue of the DOCK DHR2 domains (here, V2069, red spheres). These observations support a model, where the conserved N-terminal capping motif of DOCK10 increases its affinity for its target RhoGTPases, by rigidifying switch 2 (entropy reduction) and reinforcing contacts at the interface (enthalpy augmentation). This facilitates recruitment of the GDP-bound GTPases by the activated DOCK10, and Mg2+ expulsion by V2069 leading to the exchange process and final release of activated GTP-bound GTPases.

A

```
DOCK10_HUMAN ~~~~~~~MAGERTRRFTRSLLRP
DOCK11_HUMAN ~~~~~~~~~MAEVRKFTKRLSKP
DOCK9_HUMAN  MSQPPLLPASAETRKFTRALSKP
```

B

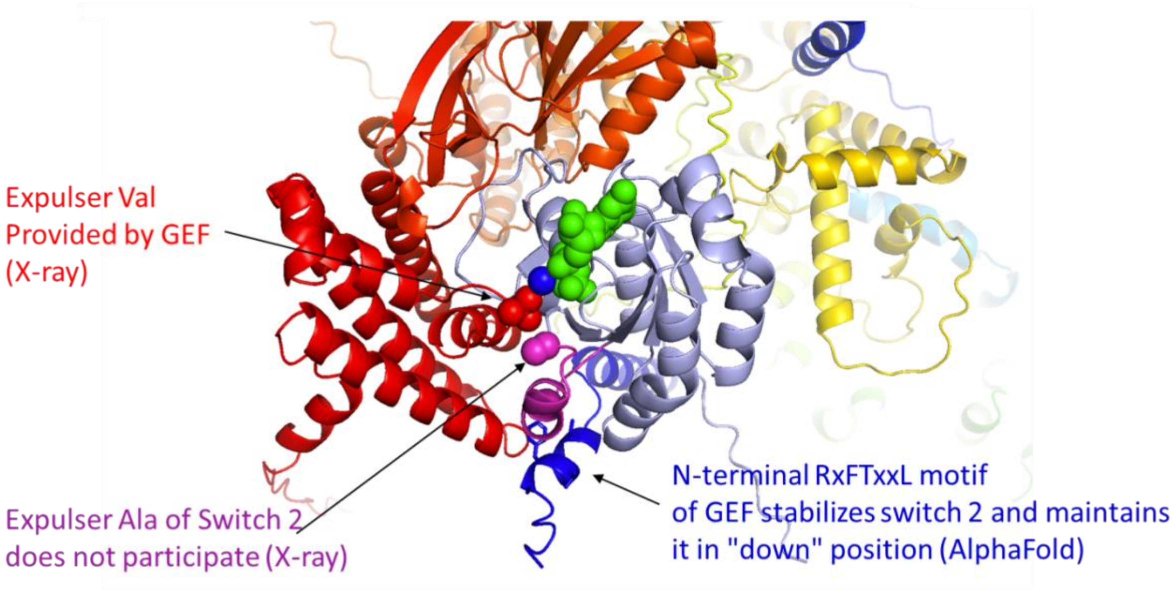



1. Chang, L., et al., *Structure of the DOCK2-ELMO1 complex provides insights into regulation of the auto-inhibited state.* Nat Commun, 2020. **11**(1): p. 3464.
2. Kukimoto-Niino, M., et al., *RhoG facilitates a conformational transition in the guanine nucleotide exchange factor complex DOCK5/ELMO1 to an open state.* J Biol Chem, 2024. **300**(7): p. 107459.
3. Yang, J., et al., *Activation of Rho GTPases by DOCK exchange factors is mediated by a nucleotide sensor.* Science, 2009. **325**(5946): p. 1398-402.
